# A multifunctional LysM effector of *Botrytis cinerea* contributes to plant infection

**DOI:** 10.1101/2022.11.05.515289

**Authors:** Mélanie Crumière, Amélie De Vallée, Christine Rascle, Shamsun Nahar, Jan A.L. van Kan, Christophe Bruel, Nathalie Poussereau, Mathias Choquer

**Author notes:** Author for correspondence: Mathias Choquer. These authors contributed equally to this work (Christophe Bruel, Nathalie Poussereau, Mathias Choquer).

## Abstract

- LysM effectors are suppressors of chitin-triggered plant immunity in biotrophic and hemibiotrophic fungi. Their role in necrotrophic fungi is unclear as these last are known to activate plant defenses and induce cell death.
- To characterize the role of the *BcLysM1* gene encoding a putative LysM effector in the necrotrophic fungus *Botrytis cinerea*, its expression was followed by transcriptional fusion and by RT-qPCR *in planta*. Two tagged-recombinant proteins were produced, and two independent deletion strains were constructed and characterized.
- *BcLysM1* is induced in the early phase of infection, and more specifically in multicellular appressoria called infection cushions. The BcLysM1 protein binds the chitin in the fungus cell wall and protects hyphae against degradation by external chitinases. It is also able to sequester chitooligosaccharides and to prevent them from inducing ROS production in A. thaliana. Using mycelium as inoculum, deletion strains show a delay in infection initiation and a default in adhesion to bean leaf surfaces.
- This study demonstrates for the first time a dual role for a LysM effector in mycelium adhesion on the plant and in host defenses suppression, both of them occurring during the asymptomatic phase of infection by a necrotrophic fungus.

## Introduction

During infection by a pathogen, plant immunity is activated by cell-membrane Pattern Recognition Receptors (PRRs). These receptors can detect the presence of Pathogen-Associated Molecular Patterns (PAMPs) and then initiate Pattern-Triggered Immunity (PTI). PTI is the first line of host defense and leads to various responses, including rapid generation of Reactive Oxygen Species (ROS), expression of defense related genes and accumulation of antimicrobial enzymes, which contribute to the basal resistance of plants (Desaki *et al*., 2018; Xiao & Kachroo, 2019). A PAMP extensively studied in plant-pathogen interactions is chitin, a N-acetyl-D-glucosamine (GlcNAc) homopolymer, that can be found in the cell wall of fungi (Kombrink *et al*., 2011; Pusztahelyi, 2018). This polymer is essential to fungal cell wall integrity and contributes to 10-20% of the cell wall dry weight (Bowman & Free, 2006). By secreting apoplastic chitin lytic enzymes during infection, plants degrade the chitin of the fungus cell wall into chitooligosaccharides (COS). These short chains of chitin are recognized by PRRs as “non-self” elements and trigger plant defenses. Plant PRRs involved in COS recognition have been identified in the plasma membrane and plasmodesmata and they all contain Lysin motif domains (LysMs) for chitin-binding (Gust *et al*., 2012; Desaki *et al*., 2018; Buendia *et al*., 2018).

LysM domains are a family of carbohydrate-binding modules with a length of approximately 50 amino acids. They are present in plants but also widely distributed in bacteria, viruses, archaea and other eukaryotes such as fungi (Buist *et al*., 2008; de Jonge & Thomma, 2009; Akcapinar *et al*., 2015). To establish infection and evade plant recognition by PRRs, pathogenic fungi secrete effector proteins among which some contain one or several LysM domain(s) (de Jonge *et al*., 2011; Rovenich *et al*., 2014; Franceschetti *et al*., 2017). Such effectors have been described in biotrophic and hemibiotrophic plant pathogenic fungi that implement a strategy allowing them to feed on living plant cells without activating host defense reactions. Two modes of action are proposed to explain the host chitin-triggered immunity suppressing capacity of biotrophic or hemibiotrophic fungi LysM effectors. Some of these proteins prevent perception by the plant by protecting the fungus cell wall from degradation by external chitinases, probably by covering and masking chitin. Mg3LysM, Mg1LysM and Mgx1LysM effectors from *Mycosphaerella graminicola*, and Vd2LysM from *Verticillium dahliae* have been described to have such role (Marshall *et al*., 2011; Kombrink *et al*., 2017; Tian *et al*., 2021). The second mechanism described prevents plant perception by binding to COS thus interfering with the PRR activation, as demonstrated for Ecp6 from *Cladosporium fulvum*, Slp1 from *Magnaporthe oryzae*, ChELP1 and ChELP2 from *Colletotrichum higginsianum*, Mg3LysM, Mg1LysM, Mgx1LysM and Vd2LysM (de Jonge *et al*., 2010; Marshall *et al*., 2011; Mentlak *et al*., 2012; Sánchez-Vallet *et al*., 2013; Lee *et al*., 2014; Takahara *et al*., 2016; Kombrink *et al*., 2017; Tian *et al*., 2021). In addition, deletion of LysM effectors encoding genes in several fungi demonstrated that they contribute to virulence (Bolton *et al*., 2008; Marshall *et al*., 2011; Mentlak *et al*., 2012; Takahara *et al*., 2016; Kombrink *et al*., 2017).

Unlike biotrophic and hemibiotrophic fungi, necrotrophic fungi secrete effector proteins and toxins to activate host defenses and to regulate plant cell death (e.g. hypersensitive response, HR), in order to kill host cells at the early steps of infection. Nevertheless, genes coding for putative LysM-effectors are present in genomes of necrotrophic fungi. Recently, the effector RsLysM was studied in the necrotrophic basidiomycete *Rhizoctonia solani*. Its encoding gene is highly induced upon sugar beet infection and the RsLysM protein suppresses chitin-triggered immunity but does not protect hyphae from enzymatic hydrolysis. This suggests that necrotrophic fungi could secrete LysM effectors to perturb chitin-triggered immunity and to establish a successful infection (Dölfors *et al*., 2019). Two other studies showed that LysM effectors may contribute to full virulence in the necrotrophic ascomycetes *Penicillium expansum* and *Lasiodiplodia theobromae* (Chen *et al*., 2020; Harishchandra *et al*., 2020). However, in these studies neither chitin binding nor analyses of host immune reaction were investigated using recombinant LysM proteins.

A previous study focusing on the necrotrophic fungus *Botrytis cinerea* revealed that *BcLysM1*, a gene encoding a LysM-domain containing protein is up-regulated in the infection cushion (IC), a multicellular appressorium structure important for secretion and pathogenicity of this fungus (de Vallée *et al*., 2019; Choquer *et al*., 2021). Moreover, the BcLysM1 protein was found to accumulate in the IC secretome (Choquer *et al*., 2021) suggesting a potential role of this protein in the early steps of infection. We therefore investigated the repertoire of LysM genes in the genome of *B. cinerea*. We found 7 *BcLysM* genes and among them, *BcLysM1* is the only one coding for a putative effector showing an induced expression during *in planta* infection. We produced the BcLysM1 recombinant protein and found that it specifically binds to chitin polysaccharide and oligomers and accumulates into the fungus cell wall. We show here that the BcLysM1 recombinant protein protects the mycelium against hydrolysis by external chitinases and can suppress the chitin-triggered immunity in a host plant. Deletion of the *BcLysM1* gene leads to a delay in symptom development on bean leaves and induces stronger host immune responses. Moreover, for the first time for a LysM effector, we demonstrated that BcLysM1 has a role in adhesion (adherence) of the fungus on plant leaves.

## Materials and Methods

### Strains and culture conditions

*Botrytis cinerea* strain B05.10 was used in this study as parental strain. B05.10 and transformed strains were maintained on malt sporulation (MS) medium (malt extract 2%, glucose 0.5%, yeast extract 0.1%, tryptone 0.1%, acid hydrolysate of casein 0.1%, ribonucleic acid 0.02%, 1.6% agar; pH5.5) supplemented with 70μg/ml hygromycin B (InvivoGen) or 70μg/ml nourseothricin (ClonNAT) when necessary. Mycelia plugs (4-mm diameter) served as inocula and cultures were incubated in the dark at 21°C. For conidia production cultures were incubated 3 days in the dark then 11 days at 21°C under continuous near-UV light. For microscopic analyses of the transcriptional strain, *B. cinerea* was cultivated in PDB^1/4^ for 6 hours for conidia, 16h for mycelia observation and 3 days for infection cushion observation. Observations were done with an Axio Observer7 LSM800 (Zeiss) confocal microscope (GFP excitation: 493 nm; emission 491 nm to 574 nm). For DNA preparations, mycelium was grown on MS medium with cellophane membrane for 3 days in the dark at 21°C. Strain LBA1126 of *Agrobacterium tumefaciens* was maintained at 28°C on LB medium without NaCl supplemented with 250μg/ml spectinomycin (Sigma) and 50μg/ml kanamycin (Duchefa).

### Gene expression analysis by RT-qPCR

For *in vitro* gene expression analysis, *B. cinerea* strains were cultivated for 3 days on MS medium. For *in planta* gene expression analysis, strains were cultivated 3 days on MS medium and then inoculated on bean leaves for infection kinetic. Vegetative mycelium or infection sites, at 0, 6, 16, 24, 30 or 48 hours post inoculation, were harvested, and immediately frozen into liquid nitrogen. Samples were then freeze dried and total RNA were extracted (RNeasy Midi Kit, Qiagen). Five micrograms of DNA-free total RNA were used as template for cDNA synthesis using the SuperScript™ IV First-Strand cDNA Synthesis kit (Thermofisher), according to manufacturer’s instructions. Quantitative PCR reactions were performed in 7900HT Fast Real-Time PCR System device (Applied Biosystems) with PowerUp™ SYBR™ Green MasterMix (Applied Biosystems) and the primers listed in Supplementary Table 1. To normalize the expression levels, *BcactA* gene was used as an internal reference for *B. cinerea* and *PvEF1α* for *P. vulgaris*. The relative mRNA amounts were calculated by the ΔΔCt method (Livak & Schmittgen, 2001) from the mean of three independent determinations of the threshold cycle (Ct), and the control sample (mycelia or no infected leaf) was used as calibrator. To measure fungal biomass, 30ng gDNA extracted from infected leaves (DNeasy PlantMini kit, Qiagen) were used for qPCR using actin specific primers for *B. cinerea* and *P. vulgaris*. Experiments were realised in biological triplicate.

### Generation of *BcLysM1* gene deletion strain and complemented strain

Two *BcLysM1* deletion mutants (*ΔBcLysM1*.*1* and *ΔBcLysM1*.*2*) and the genetic complemented strain (*ΔBcLysM1C*) were obtained by protoplasts transformation (Lalève *et al*., 2014) and *Agrobacterium*-mediated-transformation (ATMT) (Rolland *et al*., 2003) respectively. For construction of the deletion cassette, approximately 1 Kb of the 5’- and 3’-non-coding regions of *BcLysM1* gene were assembled with hygromycin resistance cassette (HPH) to replace the CDS of the *BcLysM1* gene, leading to the ΔBcLysM1.2 knock-out deletion strain. A second deletion strategy by knock-in was realized in which a GFP-Botrytis-optimized cassette (opt-GFP) (Leroch *et al*., 2011) was placed under control of the *BcLysM1* promoter leading to a GFP transcriptional fusion strain named ΔBcLysM1.1. The deletion transformants were screened on MS medium with 70μg/ml hygromycin B. Extraction of genomic DNA (DNeasy PlantMini kit, Qiagen) was performed to confirm single deletion cassette insertion by Southern blot. A vector conferring nourseothricin resistance and containing the coding region of *BcLysM1* amplified from B05.10 genomic DNA, was used for complementation of the *ΔBcLysM1*.*1* mutant. This vector was transformed into *A. tumefaciens* LBA1126 strain which was used for ATMT method with *ΔBcLysM1*.*1* strain. Diagnostic PCRs on genomic DNA were performed to verify the integration events of the selected transformants.

### Proteins extraction for BcLysM1 localization

A *BcLysM1-HA* strain expressing a tagged version of the BcLysM1 protein under the control of the *Olic* promoter was constructed, using the 3HA-tag in the N-terminal region of the protein. The strain was grown for 48h at 21°C with shaking (110rpm) in MS medium, inoculated with 2.10^5^ conidia/ml. Four protein fractions were recovered from the culture: (i) the free secreted proteins obtained from culture filtrates on 150μm membrane, precipitated with TCA and mixed with SDS-PAGE loading buffer; (ii) total proteins of mycelial culture were obtained by gently grinding the mycelium in liquid nitrogen and then mixed with 4X SDS-PAGE loading buffer. For cell wall proteins enrichment (iii), the mycelium powder was re-suspended in 5 ml of extraction buffer (50 mM Hepes pH 7; 0.25M sucrose; 10μl/ml protease inhibitor cocktail (Sigma)). Samples were mixed at 4°C for 20 min on a rotating wheal mixer. Cell wall was pelleted by centrifugation at 1000g for 5 min at 4°C. The pellet was washed thrice by repeating centrifugation in extraction buffer as above and then solubilizing in 2X SDS-PAGE loading buffer; and (iv) remaining intracellular proteins from the supernatant fraction after the 1000g centrifugation were mixed with 4X SDS-PAGE loading buffer. Blotting was performed using the Biorad Mini-Protean™ system with nitrocellulose membranes subsequently washed and blocked in TBS-Tween + 5% skimmed milk powder. Primary antibodies (mouse anti-HA H9658, Sigma) were used at dilution 1:20000 in blocking solution and anti-mouse IgG-HRP (A-9044, Sigma) at dilution 1:80000 were used for secondary antibodies. Blots were developed using ECL chemiluminescence (GE Healthcare) and visualized on ChemiDoc XRS+ (BioRad).

### Phenotypic analyses and pathogenicity assays

Fungal growth was determined by measuring daily the radial diameter of colonies on solid MS medium or minimal medium (Schumacher *et al*., 2015), following central inoculation of mycelial plugs (4-mm) excised from the edge of 3-days-old colonies. Conidiation assay was conducted as described previously (Rascle *et al*., 2018). Infection assays were performed with 1-week-old cucumber (*Cucumis sativus*) cotyledons and primary French bean (*Phaseolus vulgaris* var Saxa) leaves. The plants were inoculated with a solution of 1.5×10^3^ conidia, or 4-mm agar plugs collected from 3-days-old cultures, placed in 7.5μl GB5 medium (316 mg/L Gamborg B5, Duchefa) supplemented with 1% glucose. Infected plants were incubated at 21°C under 80% relative humidity and dark-light (16h/8h) conditions. Necrosis diameters were measured at 24, 48, 72, 96 and 168 hours post inoculation. Adhesion assay on bean leaves was performed as described previously (Pérez-Hernández *et al*., 2017) with minor modification : detached leaves were inoculated with 5-mm mycelial plugs and then incubated at 21°C under 80% relative humidity and dark-light (16h/8h) conditions during 16h. For adhesion assay on polystyrene, 6-wells microplates were inoculated with 5-mm plugs and 200μl of PDB^1/4^ were added before incubation at 21°C. Adhesion strength was measured with a 0.1N dynamometer (Eisco). For conidia adhesion, 100μl of a 1.10^4^ conidia/ml solution in PDB^1/4^ were inoculated in a 96-wells microplate. After incubation during 6h or 24h, wells were washed three times with 100μl of water. 100μl of crystal violet (0,1% in acetic acid) were added and samples were incubated 20 minutes at 21°C. Three washes were performed with water and 100μl of 90% ethanol were added to the wells. After 15 minutes of incubation at room temperature, optical density at 595nm was measured. Onion epidermis penetration was realized as described previously (Siegmund *et al*., 2013).

### Heterologous production of BcLysM1 protein in *Pichia pastoris*

For protein production in *P. pastoris*, cDNA sequence of *BcLysM1* (without signal peptide) was amplified using primers to add HIS-flag to the N-terminus and EcoRI and AvrII restriction sites for directional cloning into pPICZαA expression vector (EasySelect Pichia Expression Kit, Invitrogen). After transformation by electroporation of *P. pastoris* X-33 strain, a selected transformant was cultured for 96 hours at 28°C and 210 rpm in 500ml of BMMY medium containing 1% methanol for induction of protein production. Intracellular proteins were extracted from cells according to manufacturer’s instructions and His-tagged BcLysM1 was purified using a Ni-column (IMAC Sepharose 6 fast Flow, Cytiva) and eluted with elution buffer (20mM sodium phosphate, 0.5M NaCl, 150mM imidazole, pH 7.4). The eluted protein fractions were dialyzed against binding buffer without imidazole (20mM sodium phosphate, 0.5M NaCl, pH 7.4). Verification of protein glycosylation was realized using Pierce Glycosylation detection kit according to manufacturer’s recommendations on nitrocellulose membrane. Deglycosylation of purified protein was performed in non-denaturing conditions according to manufacturer’s protocol (Protein Deglycosylation Mix II, NEB).

### Chitin binding assay

BcLysM1 produced in *P. pastoris* was adjusted to a concentration of 5μg/ml (in 800μl) and incubated with 3mg of chitin from crab shell (Sigma), chitosan (Sigma), cellulose (Sigma), xylan (Megazyme) or *B. cinerea* cell wall fraction for 3 hours at room temperature on a rotating wheal mixer. Supernatant and pellet were separated by 13000g centrifugation for 5 minutes at 4°C. Pellets were washed three times with water and subsequent centrifugation, and then resuspended in 100μl of water. Supernatants were concentrated to approximately 100μl using VivaSpin500 columns (Sartorius, 3kDa cutoff). The two fractions were then analyzed by SDS-PAGE and Coomassie blue coloration.

For *B. cinerea* cell wall extraction, 2.10^5^ conidia were inoculated on a cellophane sheet on PDA^1/4^ medium and incubated during two days at 21°C. Mycelium was collected and frozen in liquid nitrogen. After lyophilization, 700mg of mycelium were ground to powder using a TissueLyser II (Qiagen), washed 3 times in PBS and centrifuged at 3000rpm for 10 minutes at 4°C. PBS with 1% of SDS was added to sample and incubated at 95°C for 15 minutes. After 10 minutes at 4000rpm at 4°C, pellet was washed two times with cold PBS and one time with cold water with centrifugation at 4000rpm for 10 minutes between each wash. Pellet was dried by lyophilization and then used as purified cell wall.

### Chitinase protection assay

*B. cinerea* conidia were adjusted to a concentration of 1.10^4^ conidia/ml with PDB and dispensed into a 96-well microplate in aliquots of 100μl and then incubated at 21°C for 3 hours for germination. BcLysM1 protein was added to a final concentration of 2.5μM, and after 2 hours of incubation, 50μg of chitinase from *Trichoderma viride* (Sigma) was added. All samples were further incubated for 20 hours at 21°C and then observed using a Leica DMi1 microscope with LasX software.

### Reactive oxygen species measurement

ROS production measurements were performed using leaf discs (0,4mm diameter) collected from 5-weeks-old *A. thaliana* Col0 plants. Samples were placed into a 96-well microplate, rinsed with 200μl of demineralized water every 30 minutes for 2 hours and then placed in the same volume of water overnight without shaking. Water was replaced by 100μl of fresh demineralized water and the plate was incubated for another 1 hour at room temperature. Meanwhile, mixtures of (GlcNAc)_6_ (Megazyme) and dialyzed BcLysM1 were incubated for 2 hours at room temperature. In total, 500nM of (GlcNAc)_6_ were added to trigger ROS production in the absence or presence of 1μM of effector protein in a measuring solution containing 100μM of L012 substrate (Sigma Aldrich) and 40μM horseradish peroxidase (Sigma Aldrich). Chemiluminescence measurements were taken every minute over 30 min.

### Structure and alignment of BcLysM proteins

Search of LysM genes in *B. cinerea* B05.10 genome and prediction of the protein domain organization were done using the INTERPRO database (https://www.ebi.ac.uk/interpro/). Signal peptide sequence was predicted using SIGNALP v.4.0 (Petersen *et al*., 2011). Alignments of LysM domain sequences were constructed using CLUSTAL OMEGA (http://www.ebi.ac.uk/Tools/msa/clustalo/) (Larkin *et al*., 2007).

## Results

### Seven LysM genes are identified in *B. cinerea* including the putative effector *BcLysM1*

In the genome of *B. cinerea*, seven genes, named *BcLysM1-7* (Bcin02g05630, Bcin10g06140, Bcin15g03700, Bcin10g03350, Bcin12g00930, Bcin06g00960, Bcin16g00950), contain LysM domain-encoding sequences (IPR018392). As shown in Figure **1**, analysis of their corresponding protein sequences revealed additional signatures: 1) a N-terminal signal peptide in BcLysM1, 2, 6 and 7 that suggests the secretion of these proteins through the conventional secretory pathway of *B. cinerea*; 2) a second LysM domain in BcLysM2, whose ortholog is the Ecp6 effector in *C. fulvum* (Bolton *et al*., 2008); 3) in BcLysM5, a Glycine zipper 2TM domain (also named Rick-17Da_Anti, IPR008816) and a Cyanovirin-N domain (IPR011058) with a nested LysM domain, whose ortholog is MoCVNH3 of *M. oryzae* (Koharudin *et al*., 2011, 2015) and 4) a type-1 chitin-binding domain (IPR001002) localized upstream of the LysM domain in BcLysM7. Furthermore, the LysM domains of BcLysM1, 6 & 7 contain four conserved cysteine residues that make these proteins part of the fungal-specific group of LysM proteins (Akcapinar *et al*., 2015). The other 4 BcLysM proteins do not display the four conserved cysteines and are classified as fungal/bacterial LysM (Fig. **1** and Supporting Information Fig. S1). It can be noted that the asparagine residue highly conserved in LysM effectors is not found in the two LysM domains of BcLysM2 (Supporting Information Fig. S1), although it is the ortholog of the Ecp6 gene conserved in many fungi. We mention that we characterized the *ΔBcLysM2* deleted strain, but unlike other fungi we did not find any difference compared to the parental strain (data not shown).

**Fig. 1.**
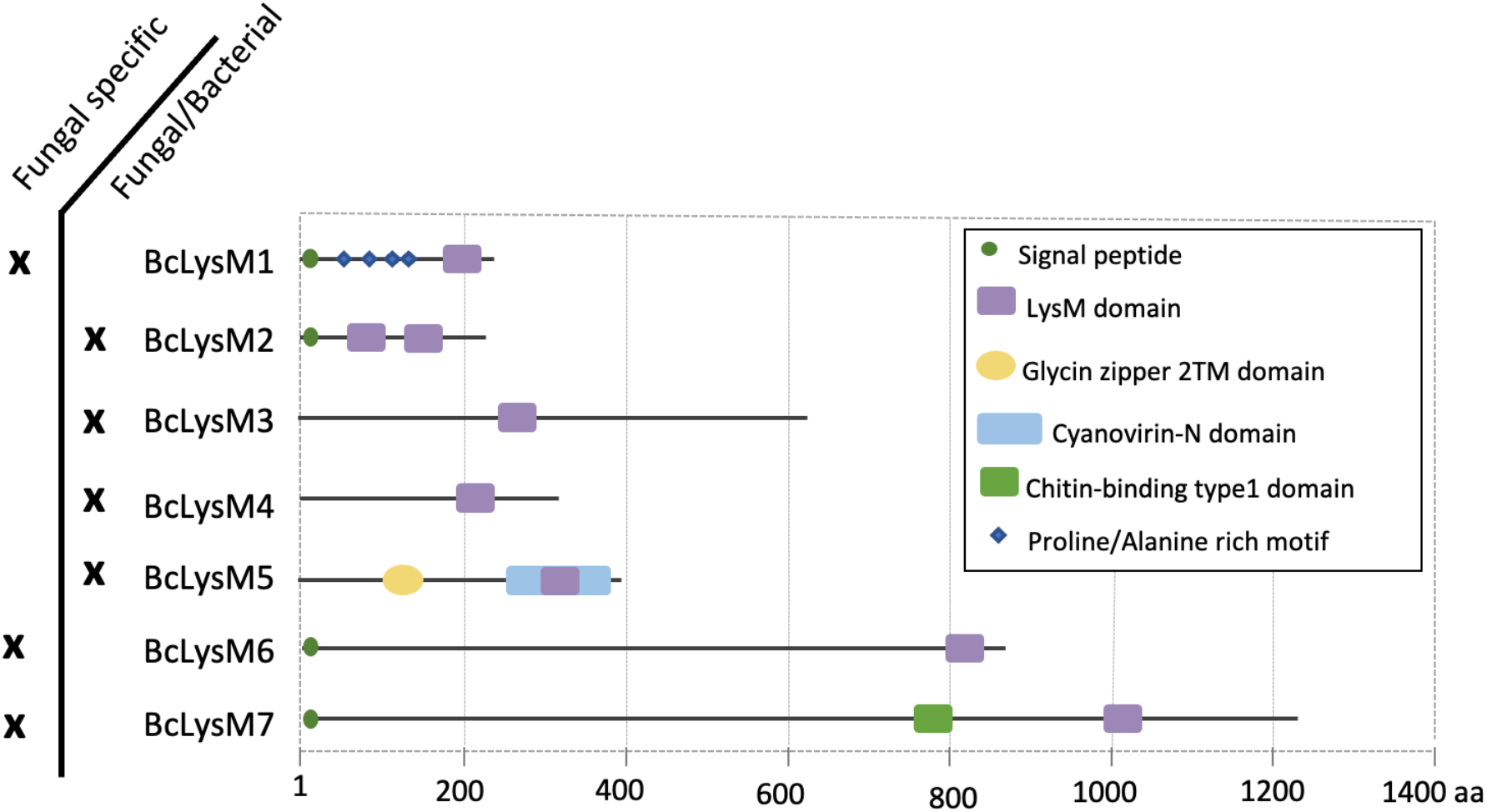
Structural features of the LysM proteins of *B. cinerea*. The genome of *B. cinerea* encodes seven putative proteins displaying 1 or 2 LysM domains detected with the Interpro domain IPR018392 and named BcLysM1 to BcLysM7. BcLysM1, BcLysM2, BcLysM6 and BcLysM7 have an N-terminal signal peptide for secretion. Four repeated Proline/Alanine rich motifs were found in the BcLysM1 sequence. The BcLysM2 protein is the only containing 2 LysM domains, BcLysM5 contains a Glycine zipper 2TM domain (IPR008816) and a Cyanovirin-N domain (IPR011058) framing the LysM domain and BcLysM7 have a chitin-binding type-1 domain (IPR001002) localized upstream of the LysM domain. The two classes of fungal LysM domains are indicated, according to Akcapinar *et al*. 2015.

Whether the *BcLysM* genes were expressed in *B. cinerea* was explored by RT-qPCR analysis of hyphae grown *in vitro* or during plant infection (Fig. **2**). All 7 genes were expressed and their expression profiles separated them in 3 groups. The first group contains *BcLysM1, 4* and *5* whose transcript levels increased (2-to 18-fold) *in planta* and peaked at 24 hours post inoculation (hpi). The second group contains *BcLysM6* whose transcript levels decreased *in planta*, and the third group contains *BcLysM2, 3* and *7* that showed similar transcript levels between both conditions. *BcLysM5* is the highest expressed LysM gene in vegetative mycelium as well as *in planta*.

**Fig. 2.**
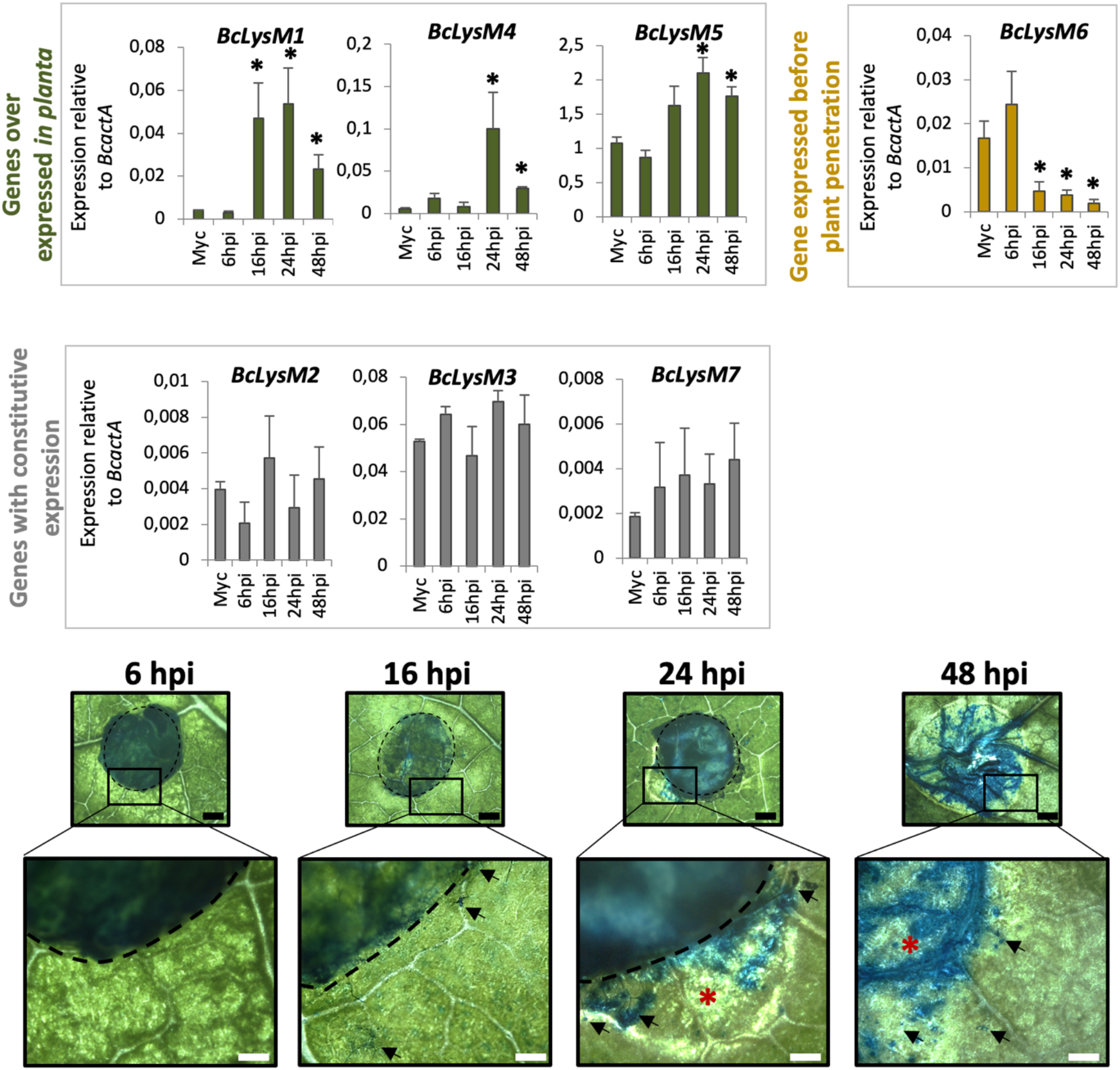
Expression of *BcLysM* genes during infection of bean leaves by *B. cinerea*. Expression was assessed using RT-qPCR and expression levels are shown relative to the *B. cinerea* reference gene *BcactA*. Data of expression levels from 6, 16, 24 and 48 hours post-inoculation (hpi) were compared to the Myc condition as control. Error bars = SE (n=3), *t*-test * = P<0.05. Kinetic of infection is showed in the lower panel, dotted black lines delimit the edge of the agar plug containing the mycelium inoculum, red stars show necrosis area visible by naked eyes and black arrows show IC. Cotton blue staining was performed to visualize the fungal cells on the leaf surface. For the 48 hpi sample, the agar plug was removed prior to taking the photograph. Black scale bars = 1mm and white scale bars = 500μm.

Among the genes induced *in planta, BcLysM1* was the only one coding for a putative secreted protein and it was chosen for further investigation.

### The *BcLysM1* gene is conserved in Sclerotiniaceae and specifically expressed in infection cushions

The *BcLysM1* gene has 25 putative orthologs in related *Sclerotiniaceae* fungi (16 other *Botrytis* spp., 4 *Monilia* spp., 3 *Sclerotinia* spp., and one *Sclerotium* spp.). The proteins share a common topology with the LysM domain located at the C-terminal end of a long region, rich in proline and alanine residues and exhibiting multiple putative N- and O-glycosylation sites (Supporting Information Fig. S2, S3). When submitting the sequence of BcLysM1 to the algorithm AlphaFold Monomer v2.0 (Jumper *et al*., 2021), a 3D model of the protein was obtained (Supporting Information Fig. S4). This model shows the LysM domain folded around two α-helices and two β-sheets and, consistent with the disruptive effect of prolines on secondary structures (Morgan & Rubenstein, 2013), the presence of “intrinsically disordered regions”.

To get information about spatio-temporal expression of the *BcLysM1* gene in *B. cinerea*, the *GFP*-encoding gene was cloned under the control of the *BcLysM1* native promoter, and the corresponding DNA cassette was introduced in *B. cinerea* (Supporting Information Fig S5a). Transformants were grown in rich liquid medium and the different stages of fungal development were observed using confocal microscopy. As shown in Figure **3**, no fluorescent signal was detected in conidia, germ tubes or undifferentiated mycelium, whereas a cytoplasmic GFP signal was detected inside mature mycelia differentiating hooks and infection cushions (IC) (Fig. **3a,b**). This result demonstrates a specific regulation of the *BcLysM1* gene and confirms its induction in IC as previously reported in a transcriptomic study (Choquer *et al*., 2021). This expression profile thus suggests a specific role for *BcLysM1* in the development and/or functioning of this fungal organ.

**Fig. 3.**
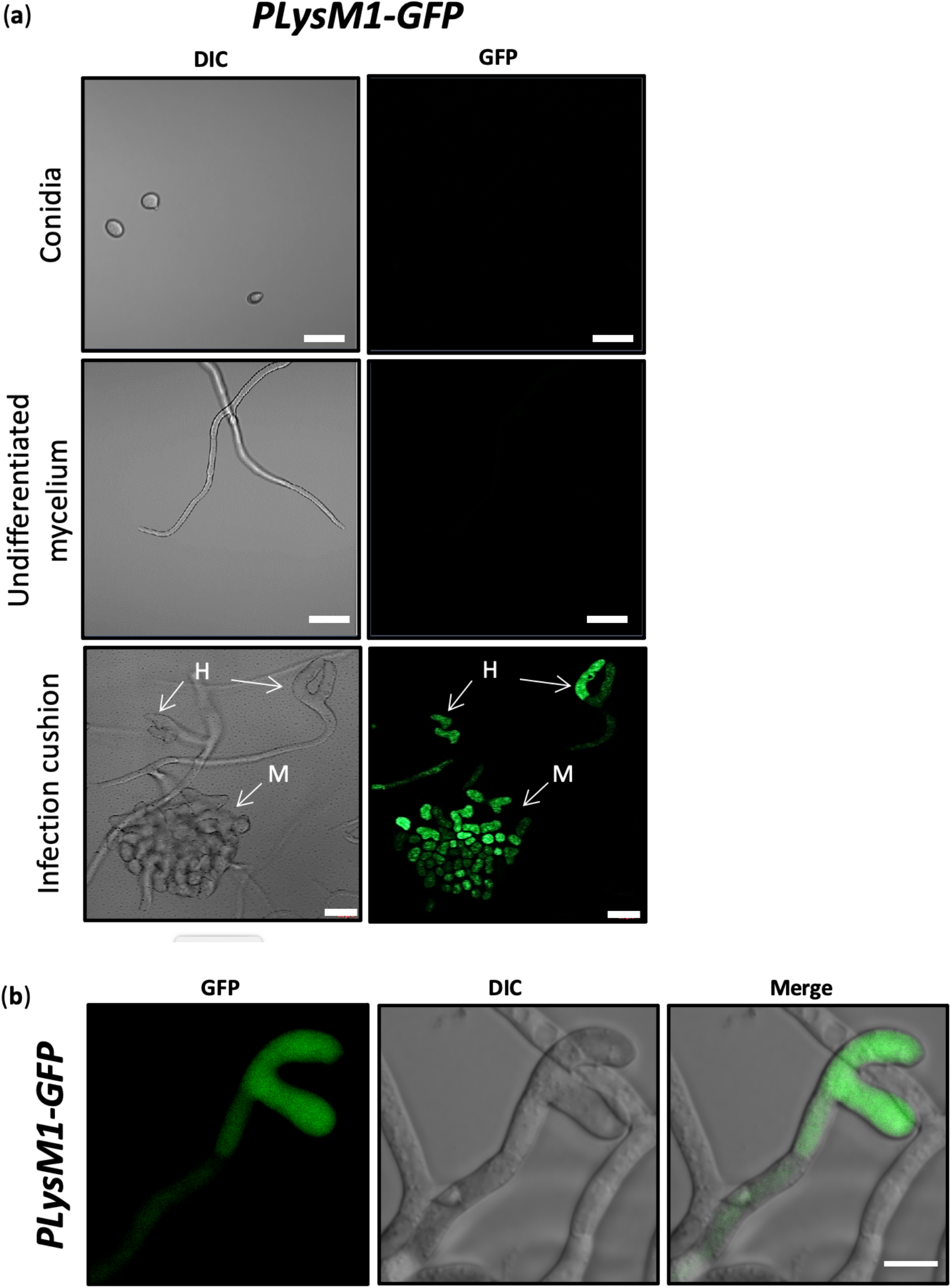
The *BcLysM1* gene expression is induced in infection cushions at hook and mature stages. (a) Confocal micrographs showing the specific expression of *P*BcLysM1-GFP transcriptional fusion, in hook stage (H) and mature stage (M) of infection cushions. No expression was observed in conidia and in undifferentiated mycelium. Scale bars = 20μm. (b) Confocal micrographs of hook stage of infection cushion. Images are maximum intensity projections from Z-stacks (ImageJ version 1.8.0), scale bar = 10 μm. DIC = Differential Interference Contrast.

### The BcLysM1 protein is hyperglycosylated, localizes in the cell wall and binds to chitin

Additional information about the BcLysM1 protein was collected by constructing a strain of *B. cinerea* expressing a *BcLysM1-3HA* encoding gene under the control of a constitutive promoter. Following growth of this transformant in liquid rich medium and preparation of mycelial crude protein extracts, the tagged BcLysM1-3HA protein could be detected at 75 kDa by using anti-HA antibodies (Fig. **4a**). Furthermore, the tagged protein was not detected among the intracellular fraction or among the secretome fraction, but it was detected among the proteins of a cell wall enriched fraction (Fig. **4a**). This result supports a mycelium cell wall localization for the BcLysM1 protein and validates its secretion by the fungal cells. In contrast, its migration as a 75 kDa protein under denaturing conditions was inconsistent with the 23 kDa predicted from its sequence. This size difference suggested post-translational modifications, as described for other LysM fungal effectors (Chen *et al*., 2014; Takahara *et al*., 2016). To explore this further, a BcLysM1-(His)_6_ recombinant protein was produced in the yeast *Pichia pastoris* and purified by metal affinity chromatography. Similarly, to BcLysM1-3HA produced in *B. cinerea*, this recombinant protein could be detected by Western Blot as a 75 kDa protein. Taking advantage of this purified protein and considering the numerous glycosylation sites present in the BcLysM1 protein sequence (Supporting Information Fig. S3), we tested the impact of N- and O-deglycosylases on the apparent molecular mass of the protein. As shown in Figure **4b**, a down-shift in migration could be monitored after enzymatic treatment, confirming the presence of sugar residues attached to BcLysM1-(His)_6_. The good efficiency (81%) of the deglycosylation reaction was verified using the periodic acid-Schiff method (Fig. **4c**) and the shift in migration was significant, yielding a protein of apparent 55-60 kDa molecular mass.

**Fig. 4.**
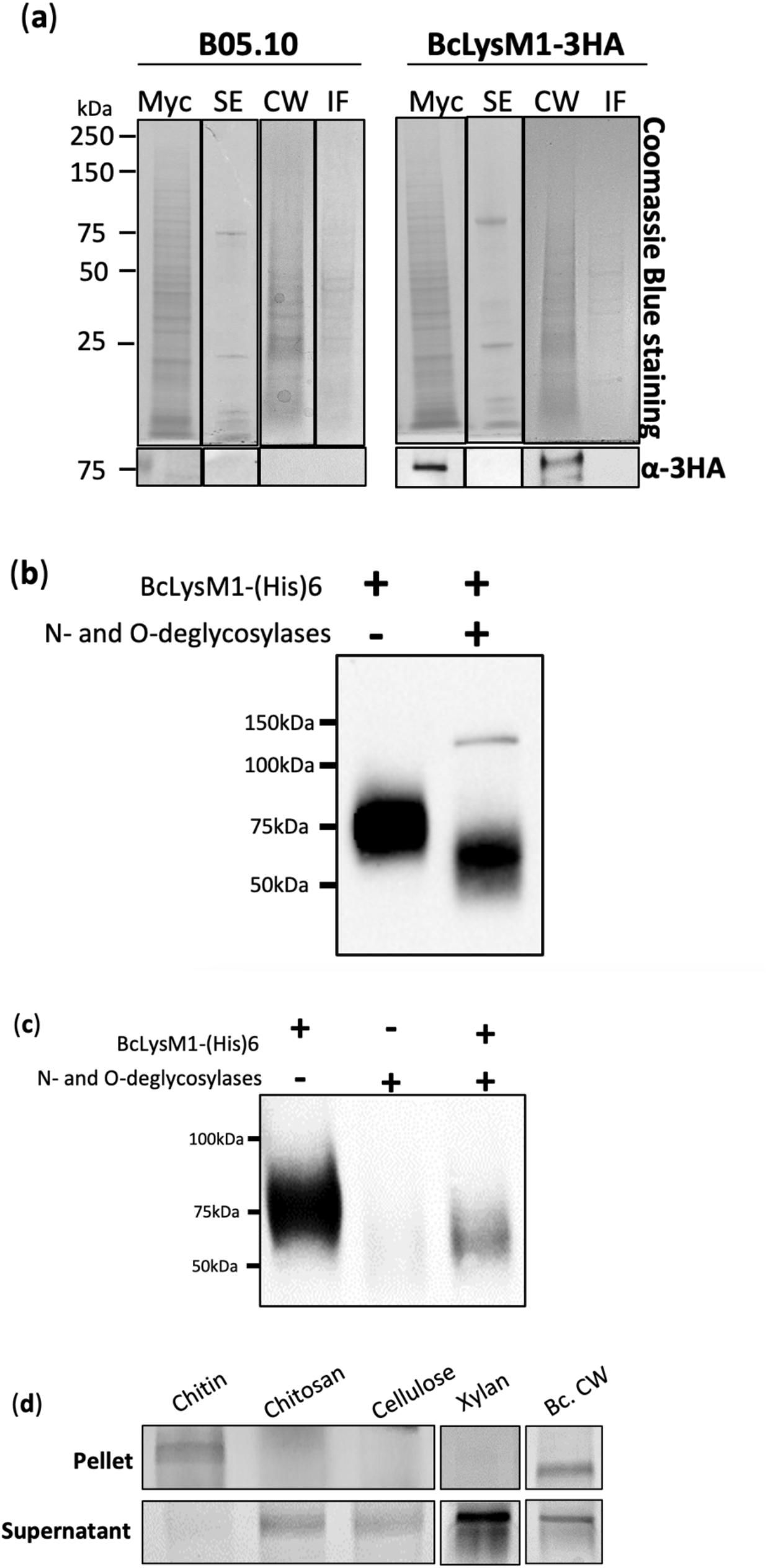
The BcLysM1 protein is highly glycosylated, binds to chitin and localizes into *B. cinerea* cell wall (a) A 3HA-tagged version of BcLysM1 expressed in *B. cinerea* B05.10 strain was used for protein detection by Western-blot with 3HA-antibody. Parental strain and BcLysM1-3HA strain were grown in liquid medium for 3 days and mycelium was recovered by filtration. A fraction of mycelium was grounded in liquid nitrogen (Myc). Differential fractionation led to separate cell wall fraction (CW) from intracellular proteins (IF). Proteins from culture secretome (SE) were also analyzed. (b) Deglycosylation of BcLysM1-(His)6 recombinant protein expressed in *P. pastoris* detected by Western-blot with anti-HIS antibody. Enzyme from the kit is also detected (at 125 kDa approximately). (c) Detection of glycosylated protein on nitrocellulose membrane using the periodic acid-Schiff method. (d) Purified BcLysM1-(His)6 protein was incubated in presence of the insoluble carbohydrates crab shell chitin, chitosan, the plant-derived carbohydrates xylan and cellulose and purified cell wall of *B. cinerea*. Following centrifugation, both the pellet and the supernatant were analyzed on protein gels and then stained by Coomassie Blue.

The purified recombinant BcLysM1-(His)_6_ was mixed with chitin, chitosan, cellulose, xylan, or purified *B. cinerea* cell wall extracts to test its binding capacity to polysaccharides (Fig. **4d**). After incubation in the presence of chitin and subsequent centrifugation, the recombinant BcLysM1 protein was detected in the chitin-containing pellet. In contrast, it remained in the supernatant when incubated in the presence of chitosan, cellulose or xylan. Lastly, it was found again in the pellet when *B. cinerea* cell wall extracts were used in the assay. These results demonstrate that BcLysM1 is a specific chitin-binding protein, likely through its LysM domain. This interaction with chitin would be sufficient to explain the observed binding of BcLysM1 to the fungal cell wall extracts. It is also consistent with the detection of BcLysM1-3HA among the cell wall proteins of the fungus mycelium (Fig. **4a**).

### *BcLysM1* deletion significantly delays mycelium-triggered infection

To investigate the biological role of BcLysM1 in *B. cinerea*, its encoding gene was deleted. Two mutant strains harboring a single insertion of the antibiotic resistance cassette at the *BcLysM1* locus were created and genetically purified to obtain homokaryons (Supporting Information Fig. S5). These two strains displayed identical phenotypes *in vitro* and *in planta* (Supporting Information Fig. S6), and one of them was selected to generate a complemented strain (Supporting Information Fig. S5c).

The *ΔBcLysM1* mutation impacted neither the radial growth rate of the fungus *in vitro*, nor the morphological appearance of the colony or the production and germination of conidia (Supporting Information Fig. S7), indicating that BcLysM1 is dispensable to the asexual saprophytic life cycle of *B. cinerea*. When bean leaves were inoculated with conidia of the parental and mutant strains, no significant difference could be observed between the two strains, indicating that BcLysM1 is also dispensable to the infection process triggered by the single appressorium produced by conidia (Fig. **5a**). Having shown that expression of the *BcLysM1* gene is induced in IC and considering that IC mostly develop from mycelium (Choquer *et al*., 2007), the inoculation assays were repeated using as inoculum agar plugs containing mycelium. In that case, a difference between the mutant and parental strains was observed. At 48 hpi, the mutant explants either produced necrosis areas smaller than the parental counterparts or triggered no visible sign of infection (Fig. **5b**). The proportion of inoculations with mutant that resulted in expanding lesions increased from 63% to 83% between 48 and 72 hpi and reached 100% at 96 hpi (Fig. **5c**), indicating a delay in infection. Once infection had begun, the expansion of the necrotic areas (colonization rate) was similar in the mutant and parental strains (Fig. **5d**). The delay in symptoms appearance was estimated at 12 hours on bean leaves (Fig. **5d**). This delay resulted in a reduced fungal biomass measured in bean leaves inoculated with the mutant strain when compared to leaves inoculated with the parental strain (−78% at 48 hpi; Fig. **5e**). In the complemented strain, the disease progression was similar to that of the wild type strain (Fig. **5b,c,d**), confirming that the *BcLysM1* deletion is responsible for the delayed-infection phenotype. The infection delay was also observed when the *ΔBcLysM1* mutant was inoculated on cucumber cotyledons (data not shown).

**Fig. 5.**
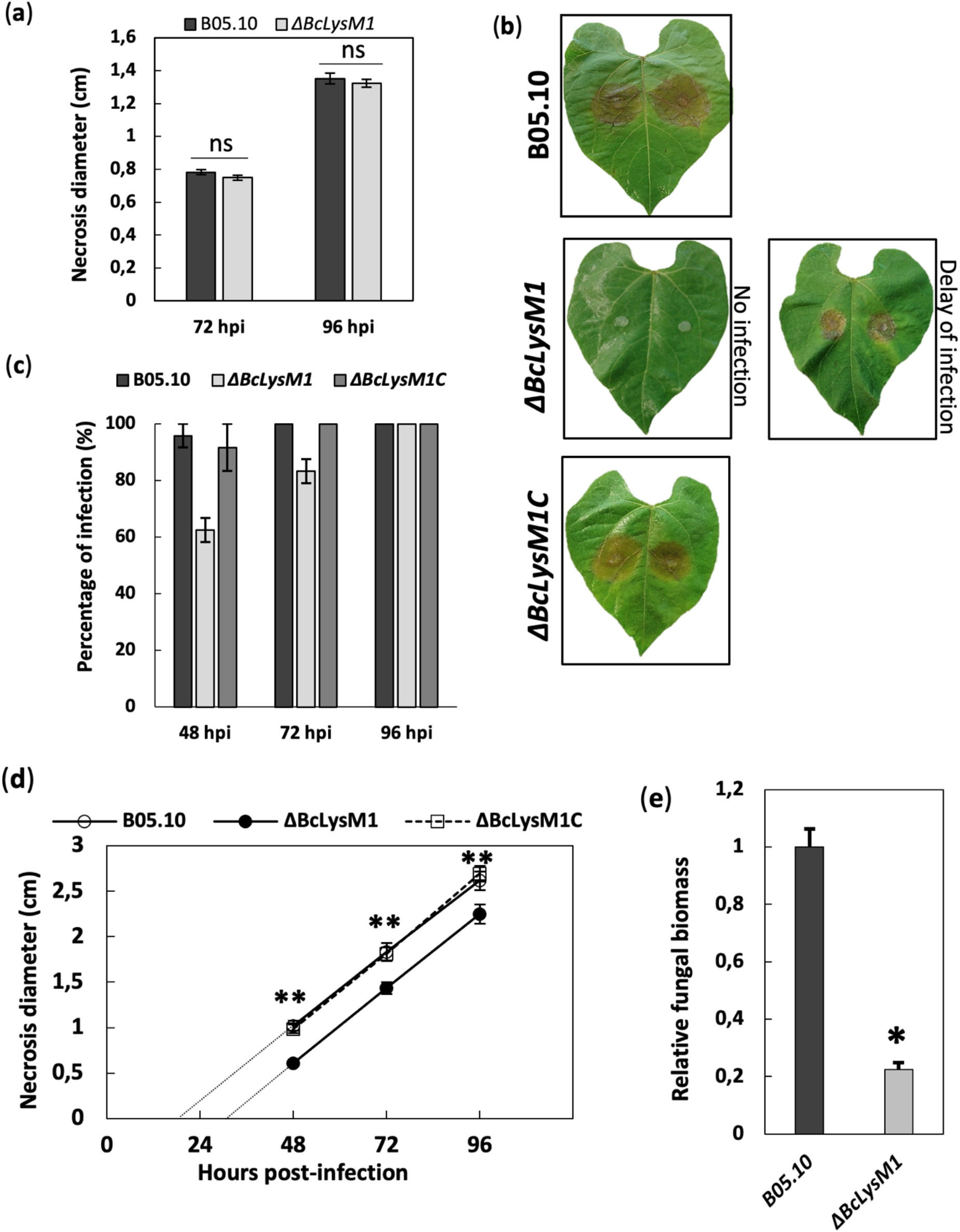
BcLysM1 deletion significantly delays mycelium-triggered infection (a) Infection assay on bean leaves with conidia inoculation (Error bars = SE, *t*-test n.s = not significant). (b) Photographs of bean leaves at 72hpi with mycelium inoculation. (c) Percentage of inoculations resulting in colonization from 2 to 4 days after inoculation on bean leaves with the B05.10 and *ΔBcLysM1* strains. (d) Measure of necrosis diameter during infection, based on the inoculation sites for which the infection had begun the second day after inoculation (Error bars = SE, *t*-test ** = P<0.01). (e) Quantification of fungal biomass during infection of bean leaves with mycelium inoculation (qPCR, quantification of *BcactA*, 48hpi, Error bars = SE, *t*-test * = P<0.05). The infection assay was realized in triplicate and the quantification was made for 2 other *Bc* genes which gave same results.

### BcLysM1 contributes to fungal adhesion on plant surface

As the initiation of leaf infection by hyphae relies on IC formation, the capacity of the *ΔBcLysM1* strain at producing these organs was investigated. The mutant mycelium grown *in vitro* or deposited on bean leaves produced IC, demonstrating that BcLysM1 is not essential for their development (Supporting Information Fig. S7). Next the capacity of the mutant IC at penetrating plant tissues was tested using onion epidermis. The similar results obtained with the mutant and parental strains indicate that BcLysM1 is not essential for IC-mediated penetration (Supporting Information Fig. S7).

As the pathogenicity assays were monitored, we noticed that some mycelial explants of the mutant could easily be detached from the bean leaves. The impact of the *ΔBcLysM1* mutation on adhesion of the fungus to the leaf surfaces was therefore explored. To this end, bean leaves were inoculated with multiple mycelial explants of mutant or parental strains and then soaked under gentle agitation. In the asymptomatic phase of infection (9 hpi; Fig. **2**) or at the appearance of symptoms (24 hpi; Fig. **2**), 72% and 91% of WT mycelial explants remained attached to the plant surface, respectively (Fig. **6a**). For the mutant strain, 55% of the mycelial explants remained attached to the plant surface at 9 hpi, and 62% at 24 hpi (Fig. **6a**; Supporting Information Fig S6). When mycelial explants were exposed to an artificial hydrophobic surface (polystyrene), a similar difference in adhesion efficacy was recorded in the mutant strain at 24 hpi (56% of attached explants for *ΔBcLysM1* and 91% for the parental strain) (Fig. **6a**). Further quantification using a dynamometer showed that the mutant mycelial explants that did attach to the bean leaves surface exhibited an adhesion strength was reduced by 50% when compared to the parental counterparts (Fig. **6b**). All these phenotypes were restored to the parental level in the complemented strain (Fig. **6a,b**). Germinated conidia of the mutant and parental strains exhibited similar adhesion capacities (Fig. **6c**). Altogether these results indicate that BcLysM1 plays a role in the adhesion capacity of a mature mycelium to the plant surface.

**Fig. 6.**
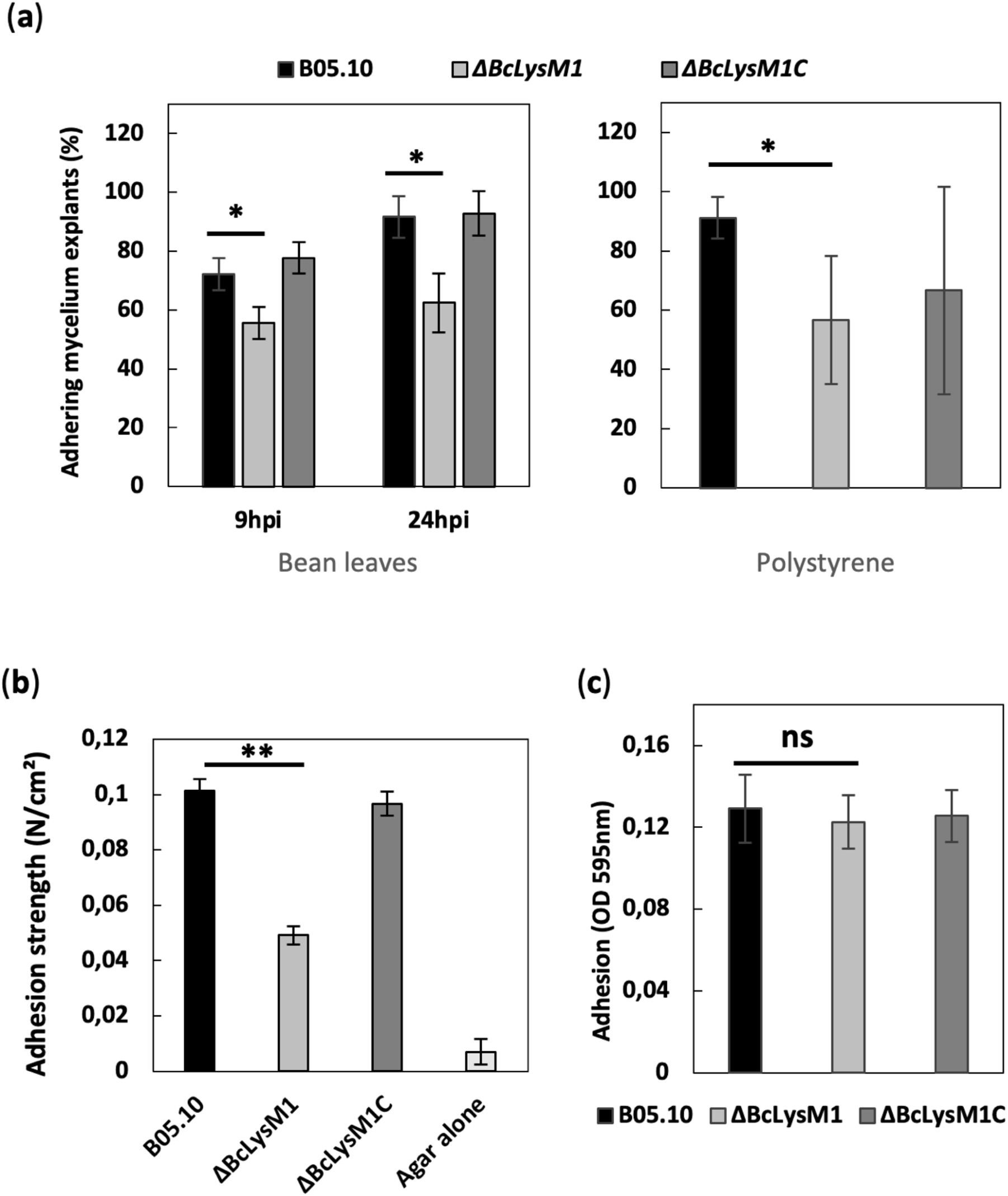
BcLysM1 contributes to mycelium adhesion to the plant surface. (a) Percentage of mycelial plugs (three days old mycelium) retained after 10 min of washing with water on bean leaves at 9 hpi and 24 hpi (n≥18) or on polystyrene surface at 24 hpi (n≥30). (b) Adhesion strength of mycelial plugs on bean leaves after 16 hpi measured with a 0.1N dynamometer (n=47). (c) Quantification of germinating conidia adhesion on hydrophobic surface. After 6 hours of incubation followed by washing and crystal violet staining, optical density at 595nm was measured. (Error bars = SE, *t*-test, *= P<0.05, **= P<0.01, ns = not significant).

### BcLysM1 protein is a suppressor of chitin-triggered plant defenses

Some of the fungal LysM effectors described in literature protect the fungal cell wall from chitinases and interfere with the activation of chitin-induced PTI in host plants (Marshall *et al*., 2011; Kombrink *et al*., 2017; Tian *et al*., 2021). Whether BcLysM1 plays such roles besides its role in fungal adhesion to host surfaces was therefore investigated. At first, conidia of the parental strain were inoculated in a rich liquid medium supplemented, or not, with chitinases from *T. viride* and with purified recombinant BcLysM1-(His)_6_. The chitinases significantly reduced germination of the conidia, but the presence of the BcLysM1 protein counteracted this deleterious effect (Fig. **7a**). As this result suggested that BcLysM1 could shield the fungal cell wall from chitinases, we reasoned that the absence of BcLysM1 in the *ΔBcLysM1* mutant should make the latter more sensitive to chitinases than the parental strain and this, indeed, was the case (Supporting Information Fig. S8a).

**Fig. 7.**
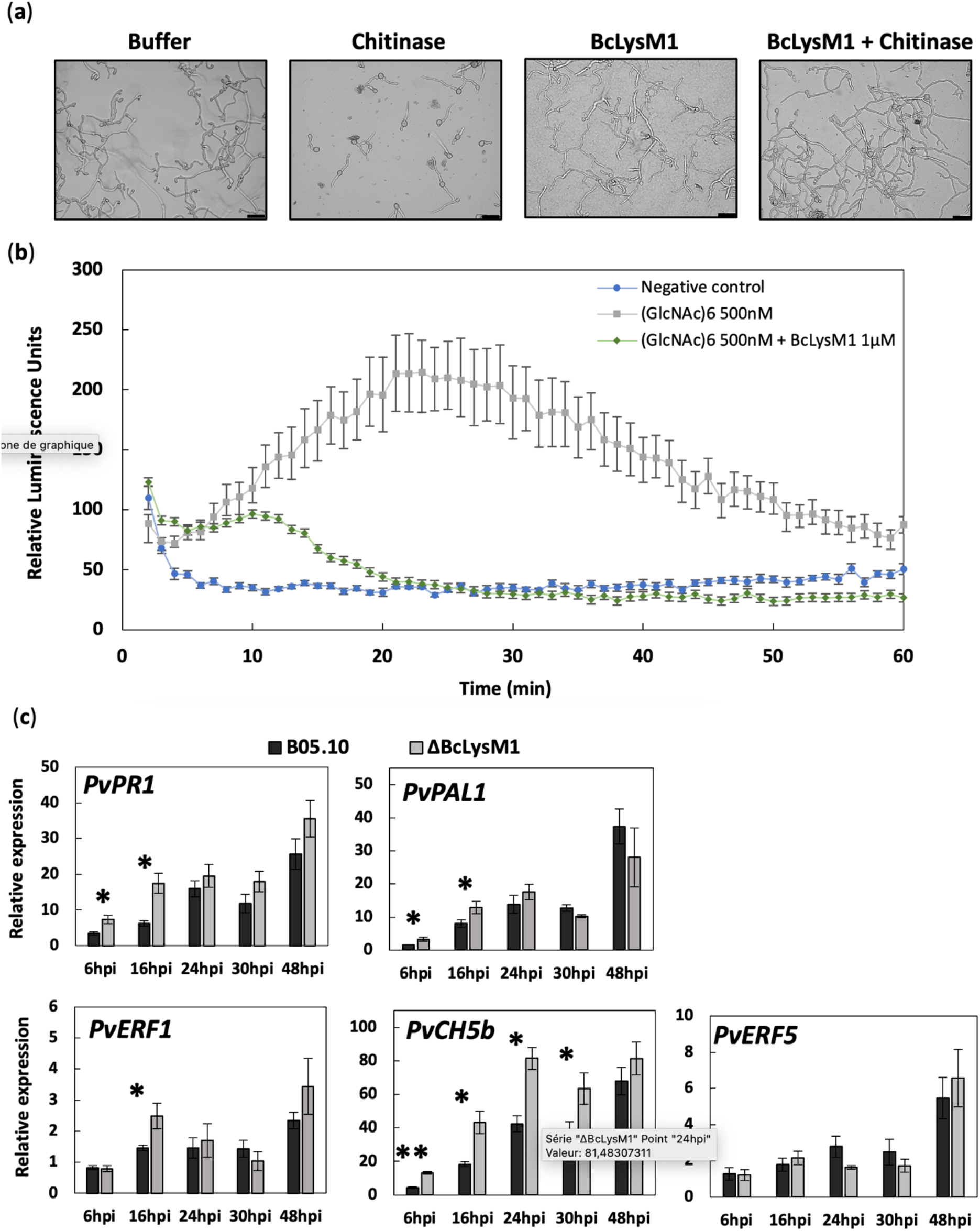
BcLysM1 protein is a suppressor of host defenses. (a) B05.10 conidia were treated with chitinases from *T. viride* with or without BcLysM1 purified protein. Photographs were taken after 20 hours of incubation. Scale bar = 50μm. (b) ROS burst assay with 4 weeks-old *A. thaliana* Col0 leaf discs. (c) Quantitative RT-PCR was performed to measure induction of several genes implicated in salicylic-acid responsive pathway (*PvPR1* and *PvPAL1*) and ethylene responsive pathway (*PvERF1, PvERF5* and *PvCH5b*). Expression data were normalized to expression of *PvEF1α* and presented relative to that detected at 0 hpi. (*t*-test, * = P<0.05 and ** = P<0.001) Error bars = SE (n=3).

Next, the capacity of the BcLysM1 protein at sequestering chitin degradation products was tested with COS (between 6 to 8 N-acetyl-glucosamine (GlNac) residues) because these products are known to induce plant defense (Liu *et al*., 2012; Gong *et al*., 2020). As shown in Figure **7b**, exposure of *A. thaliana* leaf discs to chitohexoses (6 GlNAc residues) triggered a strong production of ROS by the plant cells which, again, was counteracted by the presence of purified BcLysM1-(His)_6_. Furthermore, the sequestration of chitohexoses by BcLysM1-(His)_6_ could be demonstrated by the formation of a precipitate upon mixing of the two components *in vitro* (Supporting Information Fig. S8b). This result suggests that BcLysM1 can interfere with the detection of COS by the host plant.

To confirm that plant defenses are suppressed by BcLysM1, we investigated the expression of several genes involved in distinct plant immune response pathways. Leaves inoculated with mycelial explants of the parental and *ΔBcLysM1* mutant strains were collected over time, from the asymptomatic phase of infection (6 hpi) to host colonization and symptoms spreading (48 hpi). The samples were used to extract mRNAs and RT-qPCR was used to quantify transcriptional activation of 5 genes associated with the plant immune response (Fig. **7c**). In the leaves inoculated with the *ΔBcLysM1* mutant, the salicylic pathway-responsive genes *PvPR1* (*Pathogenesis-Related 1*) and *PvPAL1* (*Phenylalanine ammonia-lyase 1*) were 1.6 to 2.8-fold stronger induced in the early stages of infection (6 and 16 hpi) than in the leaves inoculated with the parental strain. Similarly, 2 of the 3 ethylene pathway-responsive genes analyzed, *PvERF1* (*Ethylene-Responsive transcription factor 1*) and *PvCh5b* (*Endochitinase precursor*), showed a higher expression at 16 hpi or at 6 to 30 hpi, respectively, in the leaves inoculated with the mutant strain in comparison with the parental strain (1.8-fold more induced for *PvERF1* and up to 3-fold more induced for *PvCH5b*). Expression of the third gene involved in ethylene responsive pathway, *PvERF5* (*Ethylene-Responsive transcription factor 5*), was not significantly different between infection with *ΔBcLysM1* strain or the parental strain. Taken together, these expression data indicate that BcLysM1 can reduce the detection of *B. cinerea* by a host plant and, thereby attenuate the immune response.

## Discussion

Of the 7 *BcLysM* genes found in the *B. cinerea* genome, *BcLysM1* is the only one is higher expressed during host plant infection than during mycelial growth and that encodes a secreted protein. We therefore assumed that the BsLysM1 protein could be an effector and we focused our study on this protein.

Production of a tagged version of BcLysM1 in *B. cinerea* showed that the protein is secreted and retained in the hyphal cell wall. A tagged version was also produced in *P. pastoris*, and both recombinant proteins exhibited a molecular weight higher than expected, a feature previously described for the TAL6 protein of *T. atroviride*, and for ChELP1and ChELP2 of *C. higginsianum*, produced in *P. pastoris* (Seidl-Seiboth *et al*., 2013; Takahara *et al*., 2016) and for LdpA, a LysM effector from *Aspergillus fumigatus* (Muraosa *et al*., 2019). These data suggested that post-translational modifications may occur before secretion of BcLysM1, and we indeed demonstrated that it is highly glycosylated. N-glycosylation of effectors is a mechanism used by pathogens to modulate their ability to evade host immunity (Hu *et al*., 2021; Chen *et al*., 2021). Besides, affinity of fungal LysM effectors for chitin can be mediated by glycosylation (Chen *et al*., 2014; Takahara *et al*., 2016). In *M. oryzae*, ALG3, a α-1, 3-mannosyltransferase, mediates N-glycosylation at three distinct sites in the LysM effector Slp1. Incomplete N-glycosylation of Slp1 reduces its chitin-binding capacity and thus its capacity to prevent chitin-triggered immunity in host plants (Chen *et al*., 2014). In *C. higginsianum*, the LysM proteins ChELP1 and ChELP2 are N-glycosylated and their chitin binding capacities are suppressed if the glycosyl side-chain structures are disrupted (Takahara *et al*., 2016). In *Lasiodiplodia theobromae* LtLysM1 was also described as an N-glycosylated effector protein (Harishchandra *et al*., 2020) and such glycosylation has been suggested in LysM1 and LysM2 of *Trichophyton rubrum* (Kar *et al*., 2019). In many phytopathogenic fungi, N- and O-glycosylation are involved in regulatory mechanisms associated with appressorium formation, host penetration, cell wall integrity, extracellular matrix composition and host immune evasion (Lin *et al*., 2020; Liu *et al*., 2021; Chen *et al*., 2021). In *B. cinerea*, the three O-Mannosyltransferases, PMT1, PMT2 and PMT4 are involved in the stability of the cell wall, in the production of the extracellular matrix and are required for full virulence, with a special role in adhesion and penetration into plant leaves (González *et al*., 2014). The absence of another protein, BcPMR1, a key enzyme involved in an ion transport for mannosyltransferase activity, leads to cell wall defects and impairs virulence as well as biofilm formation (Plaza *et al*., 2015). Finally, the absence of the cell wall protein BcSUN1, another protein identified in the glycoproteome of *B. cinerea* (González *et al*., 2012), reduces virulence and adhesion on plant leaves (Pérez-Hernández *et al*., 2017). In line with observations reported for many other fungal pathogens (Tronchin *et al*., 2008; Lin *et al*., 2020), these data highlight the important role of *B. cinerea* cell wall glycoproteins in adhesion.

Adhesion of a fungal cell to a host surface is the first step initiated by a pathogen to establish disease. It anchors the cell to the host surface and may also be required for host recognition and subsequent fungal development (Tucker & Talbot, 2001; Aragón *et al*., 2017). Both adhesion of conidia or mycelia have been shown critical for pathogenicity in filamentous fungi (Whiteford & Spanu, 2002; Pérez-Hernández *et al*., 2017; Fukada *et al*., 2021). In this study we show that BcLysM1 is required for mycelial adhesion to bean leaves and hydrophobic surfaces. Together with the *BcLysM1* gene being very specifically expressed in infection cushions (IC), it makes it likely that the adhesion capacity of the mycelium strongly relies on these structures. In consistence with this, BcLysM1is not involved in the adhesion of germinated conidia to surfaces and its absence does not affect bean leaf infection when conidia are used as inoculum whereas it delays infection when mycelial explants are used. Considering that germinated conidia of *B. cinerea* differentiate a unicellular appressorium to penetrate into plant while mycelium growing on plant surface differentiate ICs (Choquer *et al*., 2007), this indicates that BcLysM1 plays a significant role in mycelium-triggered infection and that adhesion of conidia-derived unicellular appressoria relies on components other than BcLysM1. As we showed that the absence of BcLysM1 does not impair the development of ICs, the role of this protein in pathogenicity relates to a subsequent step including adhesion. Interestingly, many cell wall proteins that mediate fungal adhesion share common features with BcLysM1, including high Ser/Thr content and N- or O-glycosylation (De Groot *et al*., 2005; Lipke, 2018; Tanaka & Kahmann, 2021). A role in adhesion to plant surface has not been suggested yet for a LysM effector (Tanaka & Kahmann, 2021), but the LysM1 and LysM2 proteins of *T. rubrum* have been shown to bind to N-glycoproteins of either human skin or *Neurospora crassa* cell wall (Kar *et al*., 2019).

Adhesion proteins can also prevent recognition of pathogens by host (Aimanianda & Latgé, 2010). BcLysM1 contains a single LysM domain and is able to bind chitin and protect the fungal cell wall from chitinases. Such properties, previously demonstrated for other fungal LysM proteins with a unique LysM domain (Zeng *et al*., 2020; Sánchez-Vallet *et al*., 2020; Tian *et al*., 2021), could help *B. cinerea* evading host plant recognition through chitin degradation into chitin oligomers. It has been proposed that the cell wall protection effect could be due to chitin-induced polymerisation of LysM homodimers, leading to contiguous filaments anchored to chitin in the fungal cell wall (Sánchez-Vallet *et al*., 2020; Tian *et al*., 2021). This homodimerization could be mediated by the unfolded N-terminal region of the protein (Sánchez-Vallet *et al*., 2020; Tian *et al*., 2021). Interestingly, BcLysM1 contains such region (Supporting information Fig. S3b, S7b) and precipitates in the presence of chitohexoses, suggesting its possible polymerization in the presence of chitin. BcLysM1 also contains multiple and repeated proline-rich domains, a feature described for CIH1, a LysM effector from *Colletotrichum lindemuthianum* localized in the cell wall of biotrophic hyphae during bean infection (Perfect *et al*., 1998, 2000; Takahara *et al*., 2016). The authors suggest that it could mimic and replace plant cell wall proteins at the fungal–plant interface (Perfect *et al*., 2000) and such role for BcLysM1 would further support its importance in evading plant recognition via camouflage action.

*B. cinerea* is described as a necrotrophic plant pathogen. It induces host defense to promote host cell death using cell death-inducing proteins and toxins to trigger plant cells necrosis (Bi *et al*., 2021; Leisen *et al*., 2022). However, it was demonstrated that the BcSpd1 Zn(II)_2_Cys6 transcription factor is a potential suppressor of plant defense at an early stage (14 hpi) of *A. thaliana* infection by *B. cinerea* (Chen *et al*., 2022). In this study, we show that BcLysM1 can reduce plant immune responses. This is compatible with the hypothesis of *B. cinerea* proceeding through an asymptomatic phase before triggering host cell death for colonization (Eizner *et al*., 2017; Veloso & van Kan, 2018) and it supports the importance that the suppression of the plant defenses may play during this asymptomatic phase.

They are only few studies showing that effectors of necrotrophic fungi can suppress plant immunity during infection (Shao *et al*., 2021). Our results provide a new example of this mechanism, through a LysM effector with three separate, yet complementary biological activities that are relevant for the successful infection of a host plant. As BcLysM1 has also a role in adhesion to plant host surface, we could speculate that the asymptomatic phase of *B. cinerea* infections corresponds to the time necessary for adhesion and preparation of the penetration and subsequent necrotrophic phase.

## Acknowledgements

We thank Glen Calvar and Adrien Hamandjian for their critical comments on the manuscript, Matthieu Blandenet for providing the protocol of *B. cinerea* cell wall extraction and Lisha Zhang for assistance in fungal transformation experiments. We thank Hasna Boubakri, Mathilde Fagard, Richard O’Connell and Guy Condemine for their critical advice.

## Author contributions

Original concept (MCh, NP, CB); Writing the manuscript (MCr, MCh); Editing the manuscript (MCr, MCh, NP, CB, CR, JvK, FXG); Bioinformatic analysis (MCr, MCh); Protein production (CR, MCr); Generation of *B. cinerea* mutants (AdV, SN, MCr); Phenotyping of *B. cinerea* mutants and other experiments (MCr, AdV, CR).

## Supporting Information

**Fig. S1** Alignments of the LysM domains from BcLysM proteins with LysM domains of other fungi.

**Fig. S2** 2D protein prediction alignment of BcLysM1 from *Botrytis cinerea* with its putative orthologs in other Sclerotiniaceae species.

**Fig. S3** Putative glycosylation sites od BcLysM1.

**Fig. S4** Prediction of the three-dimensional structure of BcLysM1.

**Fig. S5** Construction of the deleted and complemented strains.

**Fig. S6** Identical results obtained with a second BcLysM1 deleted strain.

**Fig. S7** BcLysM1 deletion strain is not affected in *in vitro* growth and infection cushion (IC) formation.

**Fig. S8** BcLysM1 recombinant protein protects mycelium from degradation by chitinases and binds to chitohexose.

**Table S1** Oligonucleotides used in this study.

## References

Aimanianda V, Latgé J-P. 2010. Fungal hydrophobins form a sheath preventing immune recognition of airborne conidia. Virulence 1: 185–187.

Akcapinar GB, Kappel L, Sezerman OU, Seidl-Seiboth V. 2015. Molecular diversity of LysM carbohydrate-binding motifs in fungi. Current Genetics 61: 103–113.

Aragón W, Reina-Pinto JJ, Serrano M. 2017. The intimate talk between plants and microorganisms at the leaf surface. Journal of Experimental Botany 68: 5339–5350.

Bi K, Scalschi L, Jaiswal N, Mengiste T, Fried R, Sanz AB, Arroyo J, Zhu W, Masrati G, Sharon A. 2021. The Botrytis cinerea Crh1 transglycosylase is a cytoplasmic effector triggering plant cell death and defense response. Nature Communications 12: 2166.

Bolton MD, van Esse HP, Vossen JH, de Jonge R, Stergiopoulos I, Stulemeijer IJE, van den Berg GCM, Borrás-Hidalgo O, Dekker HL, de Koster CG, et al. 2008. The novel Cladosporium fulvum lysin motif effector Ecp6 is a virulence factor with orthologues in other fungal species. Molecular Microbiology 69: 119–136.

Bowman SM, Free SJ. 2006. The structure and synthesis of the fungal cell wall. BioEssays 28: 799–808.

Buendia L, Girardin A, Wang T, Cottret L, Lefebvre B. 2018. LysM Receptor-Like Kinase and LysM Receptor-Like Protein Families: An Update on Phylogeny and Functional Characterization. Frontiers in Plant Science 9.

Buist G, Steen A, Kok J, Kuipers OP. 2008. LysM, a widely distributed protein motif for binding to (peptido)glycans. Molecular Microbiology 68: 838–847.

Chen H, He S, Zhang S, A R, Li W, Liu S. 2022. The Necrotroph Botrytis cinerea BcSpd1 Plays a Key Role in Modulating Both Fungal Pathogenic Factors and Plant Disease Development. Frontiers in Plant Science 13: 820767.

Chen D, Li G, Liu J, Wisniewski M, Droby S, Levin E, Huang S, Liu Y. 2020. Multiple transcriptomic analyses and characterization of pathogen-related core effectors and LysM family members reveal their differential roles in fungal growth and pathogenicity in Penicillium expansum. Molecular Genetics and Genomics.

Chen H, Raffaele S, Dong S. 2021. Silent control: microbial plant pathogens evade host immunity without coding sequence changes. FEMS Microbiology Reviews 45.

Chen X-L, Shi T, Yang J, Shi W, Gao X, Chen D, Xu X, Xu J-R, Talbot NJ, Peng Y-L. 2014. N - Glycosylation of Effector Proteins by an α-1,3-Mannosyltransferase Is Required for the Rice Blast Fungus to Evade Host Innate Immunity. The Plant Cell 26: 1360–1376.

Choquer M, Fournier E, Kunz C, Levis C, Pradier J-M, Simon A, Viaud M. 2007. Botrytis cinerea virulence factors: new insights into a necrotrophic and polyphageous pathogen. FEMS Microbiology Letters 277: 1–10.

Choquer M, Rascle C, Gonçalves IR, de Vallée A, Ribot C, Loisel E, Smilevski P, Ferria J, Savadogo M, Souibgui E, et al. 2021. The infection cushion of Botrytis cinerea: a fungal ‘weapon’ of plant-biomass destruction. Environmental Microbiology 23: 2293–2314.

De Groot PWJ, Ram AF, Klis FM. 2005. Features and functions of covalently linked proteins in fungal cell walls. Fungal Genetics and Biology 42: 657–675.

Desaki Y, Miyata K, Suzuki M, Shibuya N, Kaku H. 2018. Plant immunity and symbiosis signaling mediated by LysM receptors. Innate Immunity 24: 92–100.

Dölfors F, Holmquist L, Dixelius C, Tzelepis G. 2019. A LysM effector protein from the basidiomycete Rhizoctonia solani contributes to virulence through suppression of chitin-triggered immunity. Molecular Genetics and Genomics 294: 1211–1218.

Eizner E, Ronen M, Gur Y, Gavish A, Zhu W, Sharon A. 2017. Characterization of Botrytis-plant interactions using PathTrack© -an automated system for dynamic analysis of disease development. Molecular Plant Pathology 18: 503–512.

Franceschetti M, Maqbool A, Jiménez-Dalmaroni MJ, Pennington HG, Kamoun S, Banfield MJ. 2017. Effectors of Filamentous Plant Pathogens: Commonalities amid Diversity. Microbiology and Molecular Biology Reviews : MMBR 81.

Fukada F, Rössel N, Münch K, Glatter T, Kahmann R. 2021. A small Ustilago maydis effector acts as a novel adhesin for hyphal aggregation in plant tumors. The New phytologist.

Gong B-Q, Wang F-Z, Li J-F. 2020. Hide-and-Seek: Chitin-Triggered Plant Immunity and Fungal Counterstrategies. Trends in Plant Science 25: 805–816.

González M, Brito N, González C. 2012. High abundance of Serine/Threonine-rich regions predicted to be hyper-O-glycosylated in the secretory proteins coded by eight fungal genomes. BMC Microbiology 12: 213.

González M, Brito N, González C. 2014. Identification of glycoproteins secreted by wild-type Botrytis cinerea and by protein O-mannosyltransferase mutants. BMC Microbiology 14: 254.

Gust AA, Willmann R, Desaki Y, Grabherr HM, Nürnberger T. 2012. Plant LysM proteins: modules mediating symbiosis and immunity. Trends in Plant Science 17: 495–502.

Harishchandra DL, Zhang W, Li X, Chethana KWT, Hyde KD, Brooks S, Yan J, Peng J. 2020. A LysM Domain-Containing Protein LtLysM1 Is Important for Vegetative Growth and Pathogenesis in Woody Plant Pathogen Lasiodiplodia theobromae. The Plant Pathology Journal 36: 323–334.

Hu S-P, Li J-J, Dhar N, Li J-P, Chen J-Y, Jian W, Dai X-F, Yang X-Y. 2021. Lysin Motif (LysM) Proteins: Interlinking Manipulation of Plant Immunity and Fungi. International Journal of Molecular Sciences 22.

de Jonge R, Bolton MD, Thomma BP. 2011. How filamentous pathogens co-opt plants: the ins and outs of fungal effectors. Current Opinion in Plant Biology 14: 400–406.

de Jonge R, Kuroiwa H, Misumi O, Yoshida M, Ohnuma M, Fujiwara T, Yagisawa F, Hirooka S, Imoto Y, Matsushita K, et al. 2010. Conserved Fungal LysM Effector Ecp6 Prevents Chitin-Triggered Immunity in Plants. Science 329: 949–953.

de Jonge R, Thomma BPHJ. 2009. Fungal LysM effectors: extinguishers of host immunity? Trends in Microbiology 17: 151–157.

Jumper J, Evans R, Pritzel A, Green T, Figurnov M, Ronneberger O, Tunyasuvunakool K, Bates R, Žídek A, Potapenko A, et al. 2021. Highly accurate protein structure prediction with AlphaFold. Nature 596: 583–589.

Kar B, Patel P, Free SJ. 2019. Trichophyton rubrum LysM proteins bind to fungal cell wall chitin and to the N-linked oligosaccharides present on human skin glycoproteins (KA Borkovich, Ed.). PLOS ONE 14: e0215034.

Koharudin LMI, Debiec KT, Gronenborn AM. 2015. Structural Insight into Fungal Cell Wall Recognition by a CVNH Protein with a Single LysM Domain. Structure 23: 2143–2154.

Koharudin LMI, Viscomi AR, Montanini B, Kershaw MJ, Talbot NJ, Ottonello S, Gronenborn AM. 2011. Structure-Function Analysis of a CVNH-LysM Lectin Expressed during Plant Infection by the Rice Blast Fungus Magnaporthe oryzae. Structure 19: 662–674.

Kombrink A, Rovenich H, Shi-Kunne X, Rojas-Padilla E, van den Berg GCM, Domazakis E, de Jonge R, Valkenburg D-J, Sanchéz-Vallet A, Seidl M, et al. 2017. Verticillium dahliae LysM effectors differentially contribute to virulence on plant hosts. Molecular Plant Pathology 18: 596–608.

Kombrink A, Sánchez-Vallet A, Thomma BPHJ. 2011. The role of chitin detection in plant–pathogen interactions. Microbes and Infection 13: 1168–1176.

Lalève A, Gamet S, Walker A-S, Debieu D, Toquin V, Fillinger S. 2014. Site-directed mutagenesis of the P225, N230 and H272 residues of succinate dehydrogenase subunit B from Botrytis cinerea highlights different roles in enzyme activity and inhibitor binding. Environmental Microbiology 16: 2253–2266.

Larkin MA, Blackshields G, Brown NP, Chenna R, McGettigan PA, McWilliam H, Valentin F, Wallace IM, Wilm A, Lopez R, et al. 2007. Clustal W and Clustal X version 2.0. Bioinformatics 23: 2947–2948.

Lee W-S, Rudd JJ, Hammond-Kosack KE, Kanyuka K. 2014. Mycosphaerella graminicola LysM Effector-Mediated Stealth Pathogenesis Subverts Recognition Through Both CERK1 and CEBiP Homologues in Wheat. Molecular Plant-Microbe Interactions 27: 236–243.

Leisen T, Werner J, Pattar P, Safari N, Ymeri E, Sommer F, Schroda M, Suárez I, Collado IG, Scheuring D, et al. 2022. Multiple knockout mutants reveal a high redundancy of phytotoxic compounds contributing to necrotrophic pathogenesis of Botrytis cinerea. PLoS pathogens 18: e1010367.

Leroch M, Mernke D, Koppenhoefer D, Schneider P, Mosbach A, Doehlemann G, Hahn M. 2011. Living Colors in the Gray Mold Pathogen Botrytis cinerea: Codon-Optimized Genes Encoding Green Fluorescent Protein and mCherry, Which Exhibit Bright Fluorescence. Applied and Environmental Microbiology 77: 2887–2897.

Lin B, Qing X, Liao J, Zhuo K. 2020. Role of Protein Glycosylation in Host-Pathogen Interaction. Cells 9: 1022.

Lipke P. 2018. What We Do Not Know about Fungal Cell Adhesion Molecules. Journal of Fungi 4: 59.

Liu T, Liu Z, Song C, Hu Y, Han Z, She J, Fan F, Wang J, Jin C, Chang J, et al. 2012. Chitin-Induced Dimerization Activates a Plant Immune Receptor. Science 336: 1160–1164.

Liu C, Talbot NJ, Chen X-L. 2021. Protein glycosylation during infection by plant pathogenic fungi. New Phytologist 230: 1329–1335.

Livak KJ, Schmittgen TD. 2001. Analysis of Relative Gene Expression Data Using Real-Time Quantitative PCR and the 2−ΔΔCT Method. Methods 25: 402–408.

Marshall R, Kombrink A, Motteram J, Loza-Reyes E, Lucas J, Hammond-Kosack KE, Thomma BPHJ, Rudd JJ. 2011. Analysis of Two in Planta Expressed LysM Effector Homologs from the Fungus Mycosphaerella graminicola Reveals Novel Functional Properties and Varying Contributions to Virulence on Wheat. PLANT PHYSIOLOGY 156: 756–769.

Mentlak TA, Kombrink A, Shinya T, Ryder LS, Otomo I, Saitoh H, Terauchi R, Nishizawa Y, Shibuya N, Thomma BPHJ, et al. 2012. Effector-Mediated Suppression of Chitin-Triggered Immunity by Magnaporthe oryzae Is Necessary for Rice Blast Disease. The Plant Cell 24: 322–335.

Morgan AA, Rubenstein E. 2013. Proline: The Distribution, Frequency, Positioning, and Common Functional Roles of Proline and Polyproline Sequences in the Human Proteome. PLoS ONE 8: e53785.

Muraosa Y, Toyotome T, Yahiro M, Kamei K. 2019. Characterisation of novel-cell-wall LysM-domain proteins LdpA and LdpB from the human pathogenic fungus Aspergillus fumigatus. Scientific Reports 9: 3345.

Pérez-Hernández A, González M, González C, van Kan JAL, Brito N. 2017. BcSUN1, a B. cinerea SUN-Family Protein, Is Involved in Virulence. Frontiers in Microbiology 8.

Perfect SE, O’Connell RJ, Green EF, Doering-Saad C, Green JR. 1998. Expression cloning of a fungal proline-rich glycoprotein specific to the biotrophic interface formed in the Colletotrichum–bean interaction. The Plant Journal 15: 273–279.

Perfect SE, Pixton KL, O’Connell RJ, Green JR. 2000. The distribution and expression of a biotrophy-related gene, CIH1, within the genus Colletotrichum. Molecular Plant Pathology 1: 213–221.

Petersen TN, Brunak S, von Heijne G, Nielsen H. 2011. SignalP 4.0: discriminating signal peptides from transmembrane regions. Nature Methods 8: 785–786.

Plaza V, Lagües Y, Carvajal M, Pérez-García LA, Mora-Montes HM, Canessa P, Larrondo LF, Castillo L. 2015. bcpmr1 encodes a P-type Ca2+/Mn2+-ATPase mediating cell-wall integrity and virulence in the phytopathogen Botrytis cinerea. Fungal Genetics and Biology 76: 36–46.

Pusztahelyi T. 2018. Chitin and chitin-related compounds in plant–fungal interactions. Mycology 9: 189–201.

Rascle C, Dieryckx C, Dupuy JW, Muszkieta L, Souibgui E, Droux M, Bruel C, Girard V, Poussereau N. 2018. The pH regulator PacC: a host-dependent virulence factor in Botrytis cinerea. Environmental Microbiology Reports 10: 555–568.

Rolland S, Jobic C, Fèvre M, Bruel C. 2003. Agrobacterium -mediated transformation of Botrytis cinerea, simple purification of monokaryotic transformants and rapid conidia-based identification of the transfer-DNA host genomic DNA flanking sequences. Current Genetics 44: 164–171.

Rovenich H, Boshoven JC, Thomma BP. 2014. Filamentous pathogen effector functions: of pathogens, hosts and microbiomes. Current Opinion in Plant Biology 20: 96–103.

Sánchez-Vallet A, Saleem-Batcha R, Kombrink A, Hansen G, Valkenburg D-J, Thomma BP, Mesters JR. 2013. Fungal effector Ecp6 outcompetes host immune receptor for chitin binding through intrachain LysM dimerization. eLife 2: e00790.

Sánchez-Vallet A, Tian H, Rodriguez-Moreno L, Valkenburg D-J, Saleem-Batcha R, Wawra S, Kombrink A, Verhage L, de Jonge R, van Esse HP, et al. 2020. A secreted LysM effector protects fungal hyphae through chitin-dependent homodimer polymerization (H-S Guo, Ed.). PLOS Pathogens 16: e1008652.

Seidl-Seiboth V, Zach S, Frischmann A, Spadiut O, Dietzsch C, Herwig C, Ruth C, Rodler A, Jungbauer A, Kubicek CP. 2013. Spore germination of Trichoderma atroviride is inhibitedby its LysM protein TAL6. FEBS journal 280: 1226–1236.

Shao D, Smith DL, Kabbage M, Roth MG. 2021. Effectors of Plant Necrotrophic Fungi. Frontiers in Plant Science 12: 231–244.

Siegmund U, Heller J, van Kann JAL, Tudzynski P. 2013. The NADPH Oxidase Complexes in Botrytis cinerea: Evidence for a Close Association with the ER and the Tetraspanin Pls1 (HHHW Schmidt, Ed.). PLoS ONE 8: e55879.

Takahara H, Hacquard S, Kombrink A, Hughes HB, Halder V, Robin GP, Hiruma K, Neumann U, Shinya T, Kombrink E, et al. 2016. Colletotrichum higginsianum extracellular LysM proteins play dual roles in appressorial function and suppression of chitin-triggered plant immunity. New Phytologist 211: 1323–1337.

Tanaka S, Kahmann R. 2021. Cell wall–associated effectors of plant-colonizing fungi. Mycologia 113: 247–260.

Tian H, MacKenzie CI, Rodriguez-Moreno L, Berg GCM van den, Chen H, Rudd JJ, Mesters JR, Thomma BPHJ. 2021. Three LysM effectors of Zymoseptoria tritici collectively disarm chitin-triggered plant immunity. Molecular Plant Pathology n/a: 683–693.

Tronchin G, Pihet M, Lopes-Bezerra LM, Bouchara J-P. 2008. Adherence mechanisms in human pathogenic fungi. Medical Mycology 46: 749–772.

Tucker SL, Talbot NJ. 2001. Surface Attachment and Pre-Penetration Stage Development by Plant Pathogenic Fungi. Annual Review of Phytopathology 39: 385–417.

de Vallée A, Bally P, Bruel C, Chandat L, Choquer M, Dieryckx C, Dupuy JW, Kaiser S, Latorse M-P, Loisel E, et al. 2019. A Similar Secretome Disturbance as a Hallmark of Non-pathogenic Botrytis cinerea ATMT-Mutants? Frontiers in Microbiology 10: 2829.

Veloso J, van Kan JAL. 2018. Many Shades of Grey in Botrytis–Host Plant Interactions. Trends in Plant Science 23: 613–622.

Whiteford JR, Spanu PD. 2002. Hydrophobins and the interactions between fungi and plants. Molecular Plant Pathology 3: 391–400.

Xiao X, Kachroo A. 2019. Plant Defences against Fungal Attack: Perception and Signal Transduction. In: John Wiley & Sons Ltd, ed. eLS. Chichester, UK: John Wiley & Sons, Ltd, 1–8.

Zeng T, Rodriguez-Moreno L, Mansurkhodzaev A, Wang P, Berg W van den, Gasciolli V, Cottaz S, Fort S, Thomma BPHJ, Bono J-J, et al. 2020. A lysin motif effector subverts chitin-triggered immunity to facilitate arbuscular mycorrhizal symbiosis. New Phytologist 225: 448–460.

