## Supplementary material for "A multifunctional LysM effector of *Botrytis cinerea* contributes to plant infection"

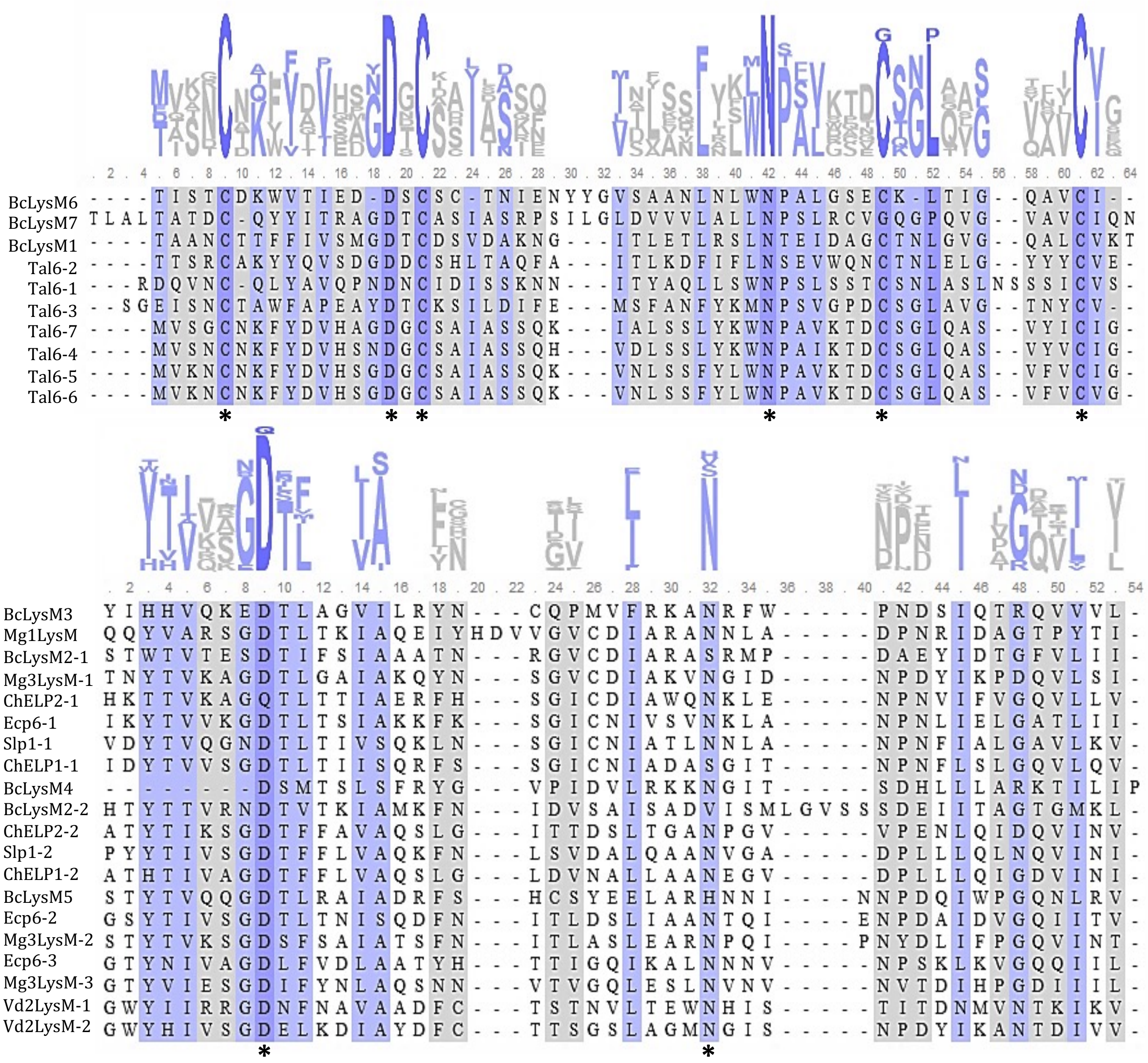


**Fig. S1** Alignments of the LysM domains from BcLysM proteins with LysM domains of other fungi. Upper panel, alignments with the fungal specific LysM domains of Tal6 from *T. atroviride*, and lower panel, alignments with the fungal/bacterial group of LysM domains (according to Ackapinar *et al.* 2015). Alignments were performed with ClustalOmega. Similarities between sequences were highlighted with dark colours. Stars show highly conserved residues in LysM domains, cysteines « C » probably implicated in two disulphide bridges, aspartic acid « D » probably implicated in ligand binding, and asparagine « N » of unknown function.

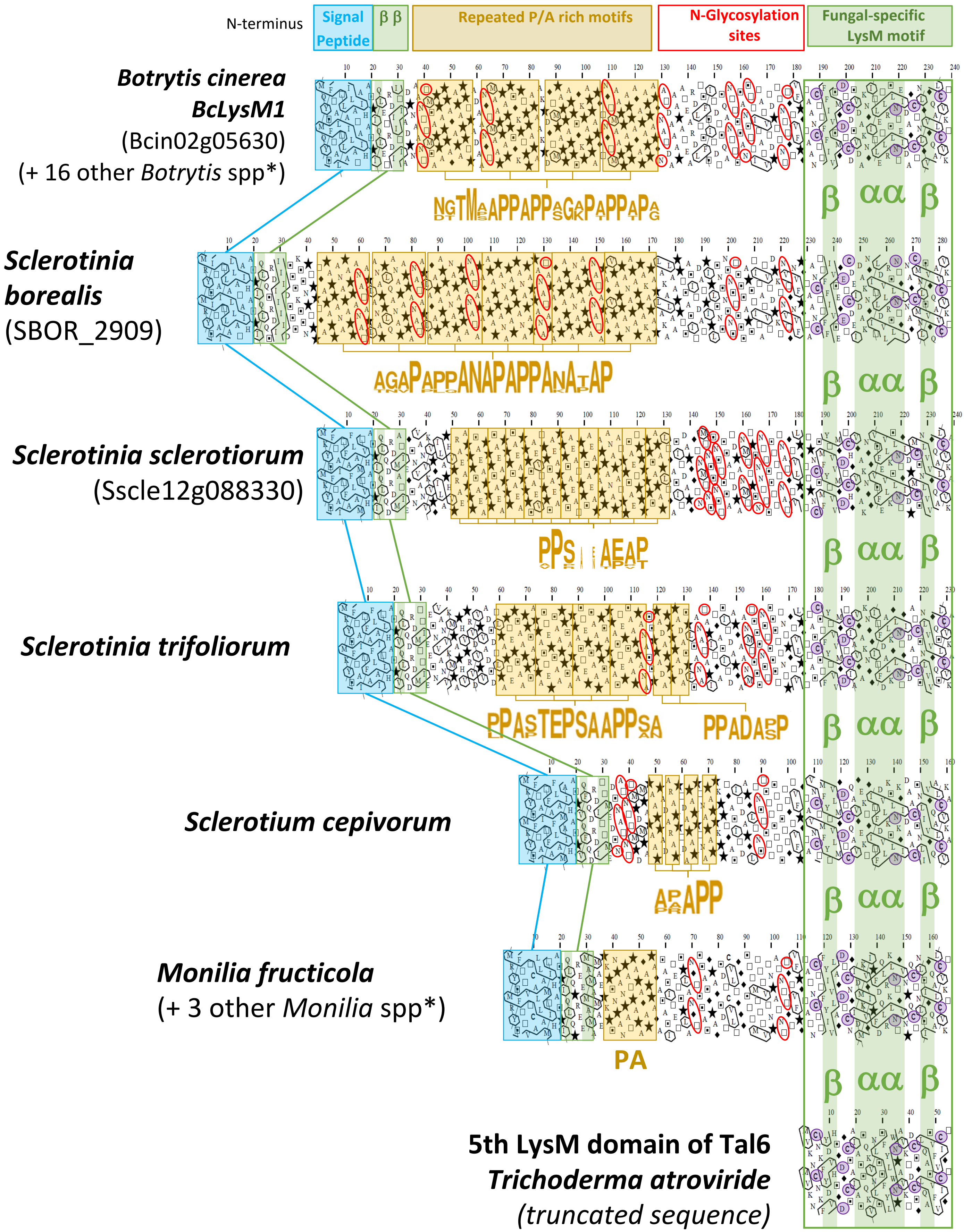
N-terminus
Signal Peptide
β β
Repeated P/A rich motifs
N-Glycosylation sites
Fungal-specific LysM motif
Botrytis cinerea
BcLysM1
(Bcin02g05630)
(+ 16 other Botrytis spp*)
Sclerotinia borealis
(SBOR_2909)
Sclerotinia sclerotiorum
(Sscle12g088330)
Sclerotinia trifoliorum
Sclerotium cepivorum
Monilia fructicola
(+ 3 other Monilia spp*)
PA
β αα β
5th LysM domain of Tal6
Trichoderma atroviride
(truncated sequence)

**Fig. S2**. 2D protein prediction alignment of BcLysM1 from *Botrytis cinerea* with its putative orthologs in other Sclerotiniaceae species. From the 1D amino acid sequences, 2D plots were created using the Hydrophobic Cluster Analysis (HCA) (http://osbornite.impmc.upmc.fr/hca/hca-form.html). On this 2D representation, the sequence is written on a duplicated α-helical net, and strong hydrophobic amino acids (V, I, L, F, M, Y, W) are encircled and their contours are joined, forming clusters (green pattern). Horizontal and vertical hydrophobic clusters are mainly associated with alpha helices and beta strands, respectively (Bitard-Feidel et al., 2018). Sequence segments separating hydrophobic clusters (at least 4 non hydrophobic amino acids or a proline) mainly correspond to loops (or hinge regions between globular domains). Symbols are used to represent amino acids with peculiar structural properties (star for proline, black diamond for glycine, square and dotted square for threonine and serine, respectively, which may be either exposed or buried). A conserved single LysM domain is found at the C-terminus extremity of LysM1 protein and its secondary structure prediction is perfectly aligned with the structural modelling of the 5th LysM domain in Tal6 protein of *Trichoderma Atroviride*, showing the typical βααβ structure of LysMs (Bayram Akcapinar *et al*., 2015). The highly conserved positions of the fungal-specific LysM consensus pattern are also found and circled in purple. The four conserved cysteine residues are probably implicated in two disulphide bridges (Cys1:Cys4 and Cys2:Cys3) stabilizing this region of the protein. The aspartic acid residue in the conserved motif GDT » is probably implicated in ligand-binding (except in *Sclerotinia borealis* where it is replaced by a glutamic acid). The role of the highly conserved asparagine residue (also found in bacteria) is unknown. At the N-terminus extremity of LysM1 proteins, a conserved signal peptide (blue box) is systematically predicted by the SignalP-5.0 Server (http://www.cbs.dtu.dk/services/SignalP/) and immediately followed by two predicted beta strands. In the centre of the LysM proteins, repeated Proline (star) and Alanine rich motifs are systematically found (yellow box and sequence logos). The role for this region of the protein is unknown but it is probably poorly structured as proline residues are known to disrupt alpha-helix or beta-sheet conformation. Finally, several N-glycosylation sites were predicted with the NetNGlyc 1.0 Server (http://www.cbs.dtu.dk/services/NetNGlyc/) and circled in red. They are mainly found between the repeated P/A rich motifs and the LysM domain and sometimes in between the P/A rich motifs as well (predicted N-glycosylation sites in the LysM domain are not shown). *A LysM1 ortholog was also found in 16 other *Botrytis* species and in 3 other *Monilia* species, displaying an identical protein topology and domain/motif composition between them (data not shown).

*MQYSFTALTLAILATAIHAS*PIQLETRDIITIADPNAA**NTTMSAPPAPPAGAPAPPTPADAANGTMAAPPAPPSGKPTPPAPAAKPD**
**GTMPAPPAPPSGKPAPPAPGTNGTMAAPPAPPAGAPAPPAPGNATDAAAPAT**EARDLTSFSITDTSQANTTSSANSTTITATVTDP
TTNTTGTVFPGTAANCTTFFIVSMGDTCDSVDAKNGITLETLRSLNTEIDAGCTNLGVGQALCVKT

**Fig. S3** Putative glycosylation sites of BcLysM1. In red O-glycosylation sites and brown N-glycosylation sites predicted by NetOGlyc and NetNGlyc server, respectively. LysM domain is underlined in black, signal peptide is represented in italics and the proline-rich region is depicted in bold.

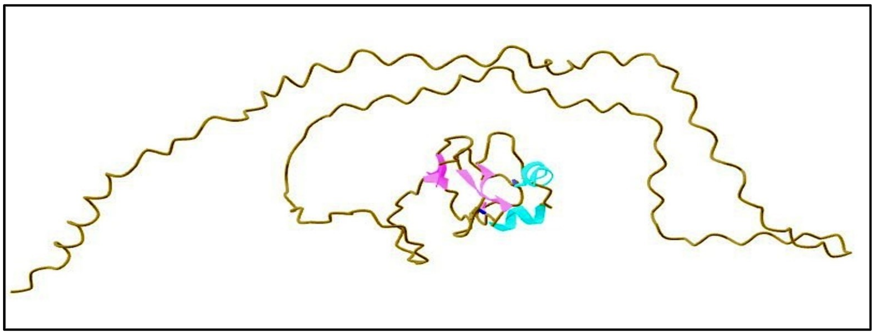

**Fig. S4** Prediction of the three-dimensional structure of BcLysM1. Prediction was realized by AlphaFold software. Beta sheet are presented in pink and alpha helix in blue.

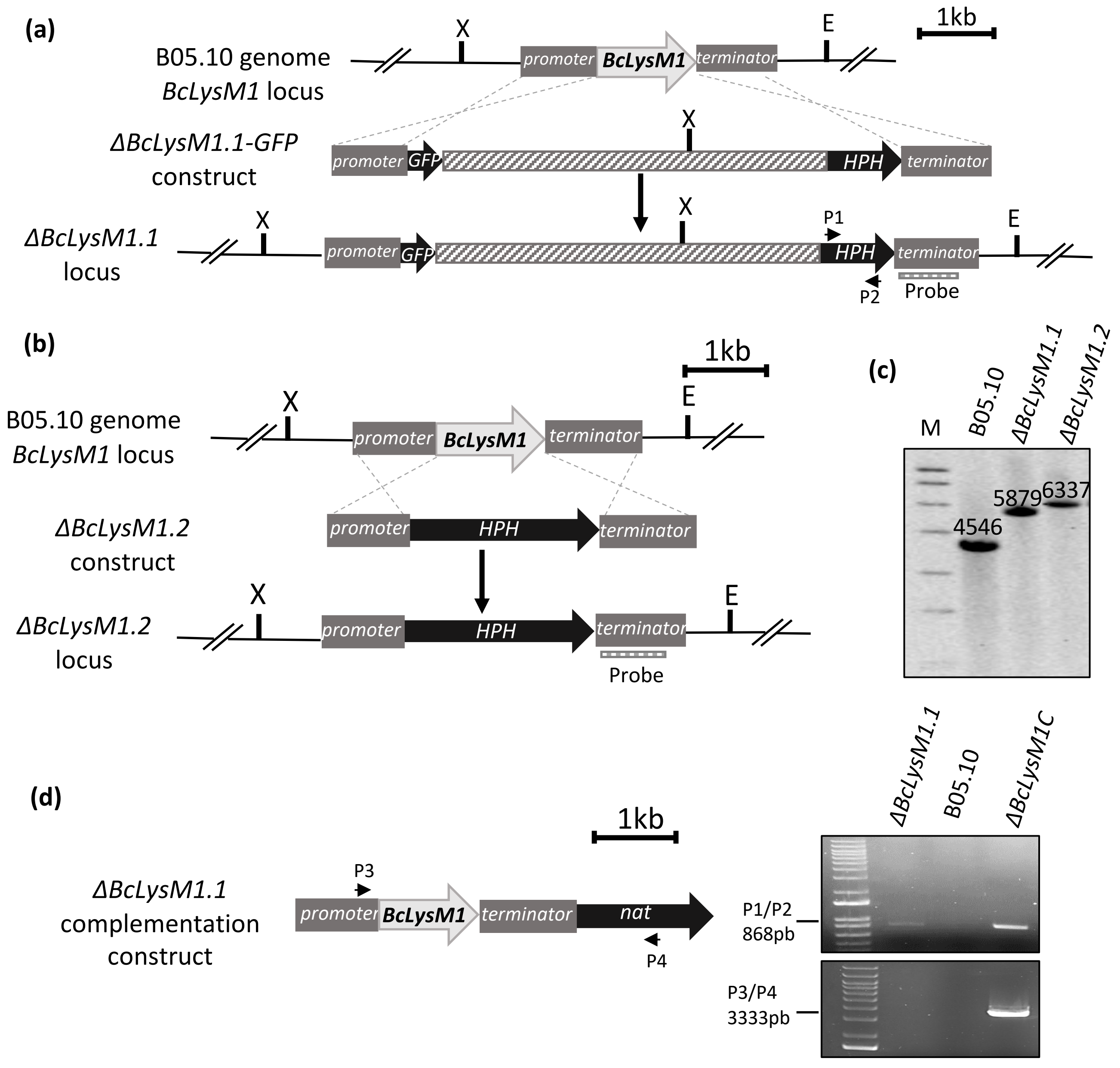


**Fig. S5.** Construction of the deleted and complemented strains. (a) Schematic representation of *BcLysM1* gene replacement by the hygromycin resistance cassette (*HPH*) flanked by 994 pb of 5’ (promoter) and 894pb of 3’ (terminator) sequences from the *BcLysM1* locus. The construction led to a transcriptional fusion by knock-in at the *BcLysM1* locus with the insertion of the GFP reporter gene (*GFP*) and the obtained strain was named *ΔBcLysM1.1* (X = XhoI, E = EcoRV). (b) Schematic representation of *BcLysM1* gene replacement by the hygromycin resistance cassette (*HPH*) flanked by 1 kb of 5’ (promoter) and 1kb of 3’ (terminator) sequences from the *BcLysM1* locus and the obtained strain was named *ΔBcLysM1.2* (X = XhoI, E = EcoRV). (c) Southern-blot analysis of the wild type strain (B05.10) and the two deleted strains (*ΔBcLysM1.1* and *ΔBcLysM1.2*). Genomic DNA was digested with EcoRV (E) and XhoI (X). Hybridization with a specific probe confirmed the presence of a unique fragment exhibiting a different size from the WT strain signal. The absence of WT signal in the two deleted strains confirmed the homokaryotic state. (M = marker). (d) Complementation construct containing the full-length of *BcLysM1* gene and a selection cassette conferring resistance to nourseothricine (nat). Diagnostic PCR to identify Δ*BcLysM1C* complemented transformants. Primers pairs are indicated.

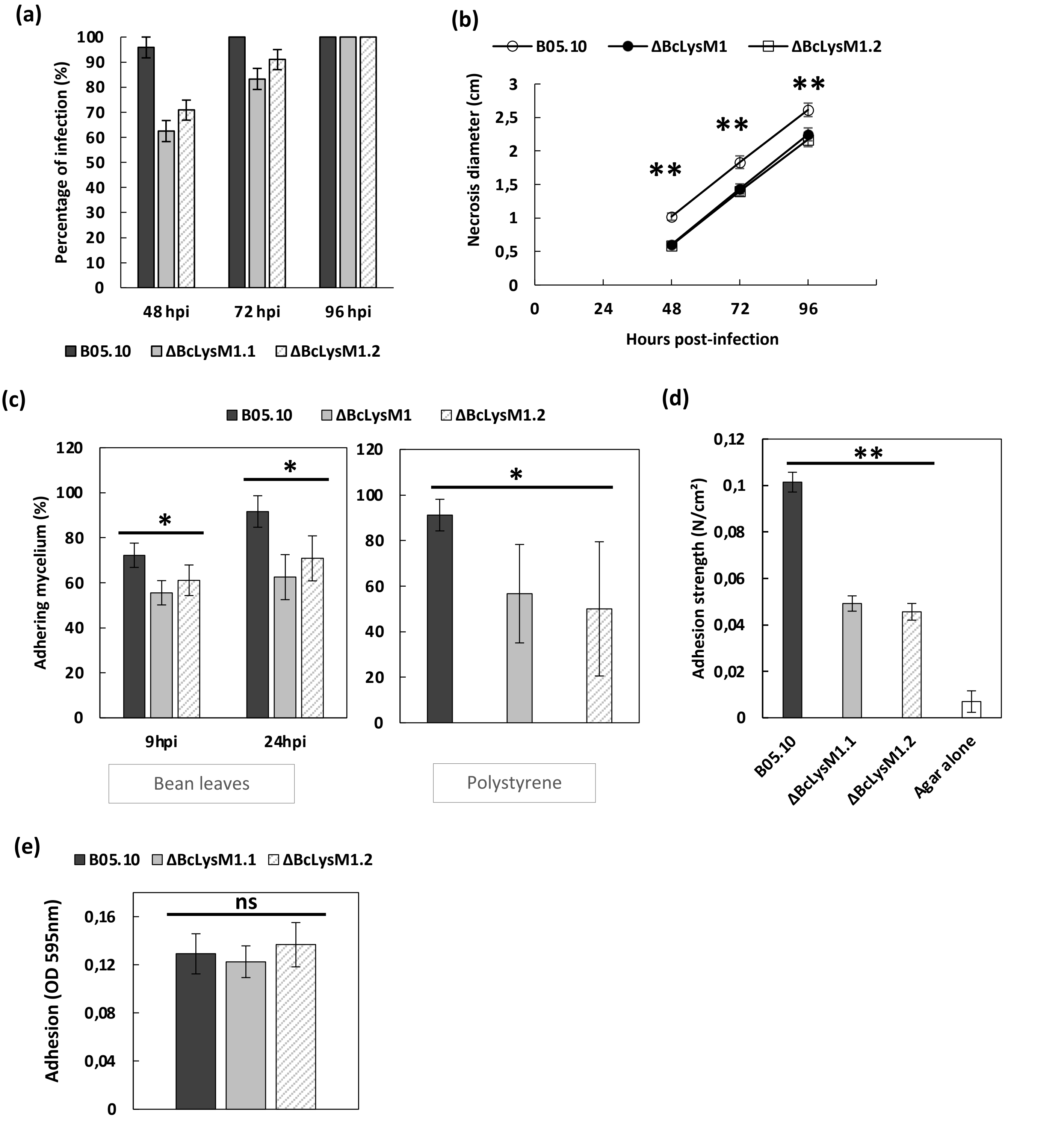


**Fig. S6.** Identical results obtained with a second BcLysM1 deleted strain. (a) Percentage of inoculations resulting in colonization from 2 to 4 days after inoculation on bean leaves with the B05.10 and *ΔBcLysM1* strains. (b) Measure of necrosis diameter during infection, based on the inoculation sites for which the infection had begun the second day after inoculation (Error bars = SE, t-test ** = $P<0.01$). (c) Percentage of mycelial plugs (three days old mycelium) retained after 10 min of washing with water on bean leaves at 9 hpi and 24 hpi ($n\geq18$) or on polystyrene surface at 24 hpi ($n\geq30$). (d) Adhesion strength of mycelial plugs on bean leaves after 16 hpi measured with a 0.1N dynamometer (n=47). (e) Quantification of germinating conidia adhesion on hydrophobic surface. After 6 hours of incubation followed by washing and crystal violet staining, optical density at 595nm was measured. (*t*-test, *= $P<0.05$, **= $P<0.01$, ns = not significant).

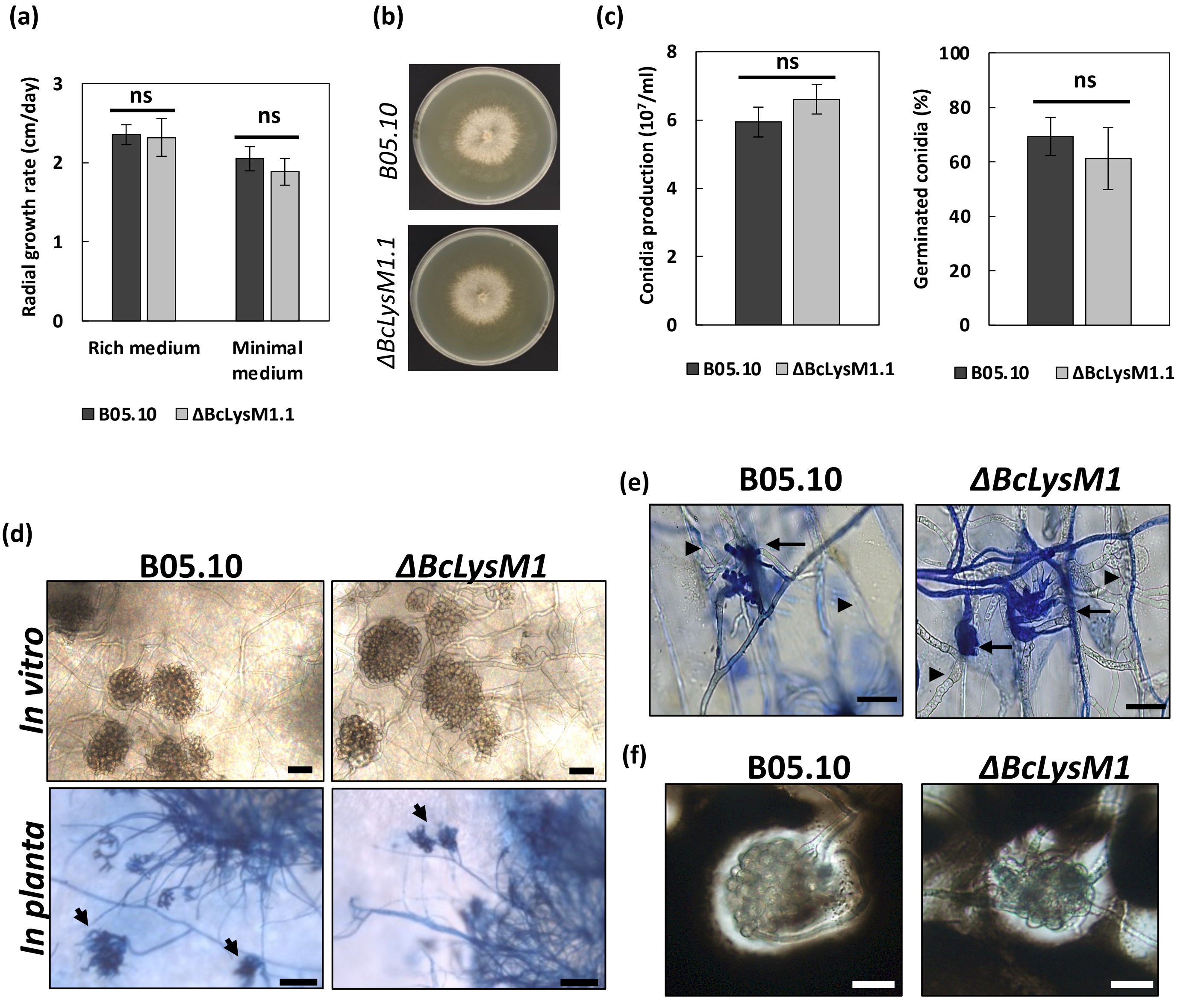


**Fig. S7.** BcLysM1 deletion strain is not affected in *in vitro* growth and infection cushion (IC) formation. (a) Growth rate on rich and minimal media. Strains were incubated at 21°C for 7 days and colony diameter was monitored each day. (b) Colony morphology after 3 days at 21°C on MS medium. (c) Left panel; conidia production on MS medium after 11 days under near-UV light. Right panel; percentage of germinated conidia on $PDB^{1/4}$ medium after 3 hours. (d) Differentiation of IC *in vitro* (top) ($PDB^{1/4}$, 21°C for 48 hours, scale bars = 50µm) and *in planta* by trypan blue coloration (bottom) (bean leave, 21°C for 48 hours, scale bars = 50µm), arrows indicate IC. (e) Penetration of infection hyphae into onion epidermis was observed at 16 hpi. Fungal cell wall was colored by cotton blue prior visualization. Arrow heads indicate infectious hyphae invading the plant (not stained), and arrows indicate epiphyte hyphae and IC on epidermis surface (blue colored). (Scale bars = 50µm). (f) China ink staining for detection of the extracellular matrix surrounding the IC (21°C for 72 hours, scale bars = 50µm).

**(a)**

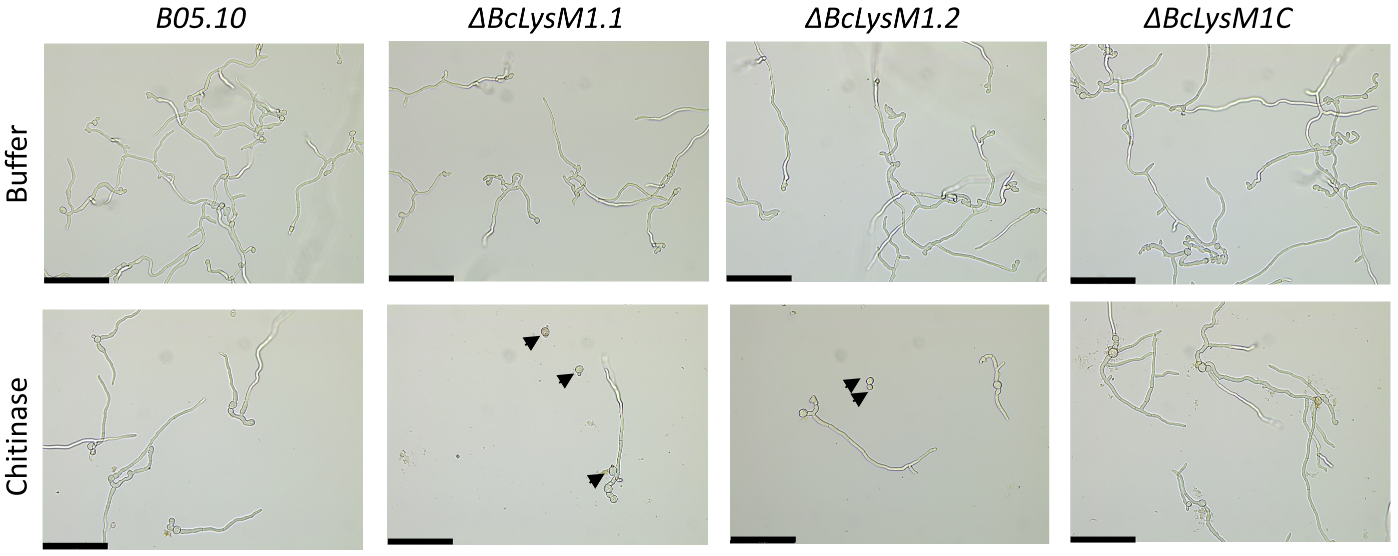


**(b)**

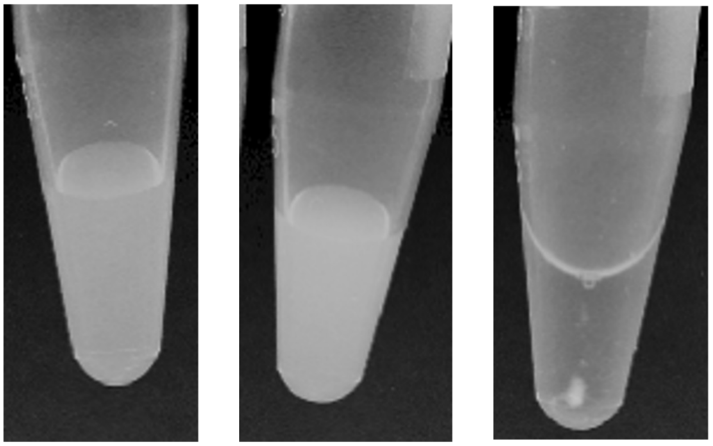


**Fig. S8** BcLysM1 protein protects mycelium from degradation by chitinases and binds to chitohexose. (a) BcLysM1 protects cell wall against chitinases. Conidia of *B. cinerea* were incubated with chitinases from *T. viride*. Photographs were taken after 18 hours of incubation. Arrows show ungerminated conidia. Scale bars = 120µm. (b) BcLysM1 binds to COS. Purified BcLysM1 produced in *P. pastoris* (40µg) was incubated with chitohexose (125nM) or water as control, at room temperature. After overnight incubation, 2µl of 0.2% methylene blue (Sigma-Aldrich) was added and incubated for 30 minutes. Samples were centrifuged at 20,000g for 15 minutes. (According to Sánchez-Vallet *et al*. 2020 and Tian *et al*. 2021).

**Table S1** Oligonucleotides used in this study.

| Gene ID | Organism | Name | Forward | Reverse | Experiment | Reference |
|---|---|---|---|---|---|---|
| Bcin02g05630 | *B. cinerea* | BcLysM1 | GATAGTGTTGACGCGAAGAATGG | AGCATCAATTTCGGTATTCAAACTC | RT-qPCR | This study |
| Bcin10g06140 | *B. cinerea* | BcLysM2 | GCAGAATGCCAGATGCTGAA | CGGCGGGAATGATCAACA | RT-qPCR | This study |
| Bcin15g03700 | *B. cinerea* | BcLysM3 | GCGATTGAAGGTTGGGTTAGAA | CGTACCGCCTCCATTTGG | RT-qPCR | This study |
| Bcin10g03350 | *B. cinerea* | BcLysM4 | TGAATACGACATTGCAGTCCTGTA | TCCCATTGCGAGCTCTAGGT | RT-qPCR | This study |
| Bcin12g00930 | *B. cinerea* | BcLysM5 | GGCGGCTCGACATACACTGT | GAAGCGGTCGGCAATAGCT | RT-qPCR | This study |
| Bcin06g00960 | *B. cinerea* | BcLysM6 | GGAAGCGAGTGTAAGTTGACCAT | CGCTTGTAGGCTCCCTTGAC | RT-qPCR | This study |
| Bcin16g00950 | *B. cinerea* | BcLysM7 | CTTCCGCTAGCGCTTGTAGAC | TGCGGTTAAGGCGAGAGTTAC | RT-qPCR | This study |
| Bcin02g05630 | *B. cinerea* | BcLysM1-prom | AACCTTAGCGAGAGATTTGAAGC | TTTGAACTTATGGATTGAATATCG | Deletion | This study |
| Bcin02g05630 | *B. cinerea* | BcLysM1-term | CAATAAAGAACGGTGACTTCTGC | AGTCTAGAGACAACTTGAAGCTCC | Deletion | This study |
| Bcin02g05630 | *B. cinerea* | BcLysM1-termProbe | GGGACAGTATAGATGCTGTACTTCG | TCAATGAGTCCTGGCTATCTCG | Southern blot | This study |
| Bcin02g05630 | *B. cinerea* | BcLysM1-com | CGGAAGGGATTTTCTTGTATTC | AGGTTGAGTCTTGGGTCG | Gene complementation | This study |
| - | - | P1 | TTTCAGCTTCGATGTAGGAGG | - | Verification complemented strain | This study |
| - | - | P2 | - | GAGTACTTCTACACAGCCATCG | | |
| - | - | P3 | TCTGTACCTCACCTCACCTG | - | | |
| - | - | P4 | - | TGCGTTGACGTTGGTGACCT | | |
| Bcin02g05630 | *B. cinerea* | BcLysM1-prodHIS | CAACCAAAGAATTCCATCATCACCATCACCACTCTCCAATTCAATTGGAGACGAGAG | GTTGGTTTCCTAGGCTAAGTCTTAACACAAAGCGCTTGTC | Protein production in *P. pastoris* | This study |
| Bcin16g02020 | *B. cinerea* | BcactA | CCGTGCTCCAGAAGCTTTGT | GTGGATACCACCGCTCTCAAG | RT-qPCR/qPCR | Rascle *et al*. 2018 |
| Phvul.008G011000 | *P. vulgaris* | PvACT | TGCATACGTTGGTGATGAGG | AGCCTTGGGGTTAAGAGGAG | RT-qPCR/qPCR | Mayo *et al*. 2016 |
| Phvul.004G060000 | *P. vulgaris* | PvEF1α | GGTCATTGGTCATGTCGACTCTGG | GCACCCAGGCATACTTGAATGACC | RT-qPCR | Guerrero-Gonzalez *et al.* 2011 |
| Phvul.003G109100 | *P. vulgaris* | PvPR1 | TGGTCCTAACGGAGGATCAC | TGGCTTTTCCAGCTTTGAGT | RT-qPCR | Mayo et al. 2016 |
| Phvul.001G177800 | *P. vulgaris* | PvPAL1 | TGAGAGAGGAGTTGGGCACT | TTCCACTCTCCAAGGCATTC | RT-qPCR | Mayo et al. 2016 |
| Phvul.009G116500 | *P. vulgaris* | PvCH5b | CAGCCAAAGGCTTCTACACC | TTGTTTCGTGAGACGTTTGC | RT-qPCR | Mayo et al. 2016 |
| Phvul.007G127800 | *P. vulgaris* | PvERF1 | CGCTCTCAAGAGGAAACACTCC | TGAATCAGAAGGAGGAGGGAAT | RT-qPCR | Mayo et al. 2016 |
| Phvul.002G055700 | *P. vulgaris* | PvERF5 | GGCTCCAAGTGGATTGAGAAC | TCAGAATCAGATAACTACAAAGCACAA | RT-qPCR | Mayo et al. 2016 |
